# Novel risk genes and mechanisms implicated by exome sequencing of 2,572 individuals with pulmonary arterial hypertension

**DOI:** 10.1101/550327

**Authors:** Na Zhu, Michael W. Pauciulo, Carrie L. Welch, Katie A. Lutz, Anna W. Coleman, Claudia Gonzaga-Jauregui, Jiayao Wang, Joseph M. Grimes, Lisa J. Martin, Hua He, PAH Biobank, Yufeng Shen, Wendy K. Chung, William C. Nichols

**Affiliations:** Department of Pediatrics, Columbia University Medical Center, New York, New York, USA; Department of Systems Biology, Columbia University, New York, New York, USA; Division of Human Genetics, Cincinnati Children’s Hospital Medical Center, Cincinnati, OH, USA; Department of Pediatrics, University of Cincinnati College of Medicine, Cincinnati, OH, USA; Regeneron Genetics Center, Regeneron Pharmaceuticals Tarrytown, New York, USA; Department of Biomedical Informatics, Columbia University, New York, New York, USA; Herbert Irving Comprehensive Cancer Center, Columbia University Medical Center, New York, New York, USA; Department of Medicine, Columbia University Medical Center, New York, New York, USA

## Abstract

Group 1 pulmonary arterial hypertension (PAH) is a rare disease with high mortality despite recent therapeutic advances. Pathogenic remodeling of pulmonary arterioles leads to increased pulmonary pressures, right ventricular hypertrophy and heart failure. Mutations in bone morphogenetic protein receptor type 2 and other risk genes predispose to disease, but the vast majority of non-familial cases remain genetically undefined. To identify new risk genes, we performed exome sequencing in a large cohort from the National Biological Sample and Data Repository for PAH. By statistical association of rare deleterious variants, we found tissue kallikrein 1 and gamma glutamyl carboxylase as new candidate risk genes for idiopathic PAH associated with a later age-of-onset and relatively moderate disease phenotype compared to bone morphogenetic receptor type 2. Both genes play important roles in vascular hemodynamics and inflammation but have not been implicated in PAH previously. These data suggest new genes, pathogenic mechanisms and therapeutic targets for this lethal vasculopathy.

## Introduction

Pulmonary arterial hypertension (PAH) is a progressive vascular disease characterized by proliferative remodeling, increased pulmonary pressures and resistance, and high mortality ^1–4^. The disease is etiologically heterogeneous, classified as familial (FPAH) as a subset of heritable PAH, idiopathic (IPAH), associated with other medical conditions (APAH, including autoimmune connective tissue disorders (CTD), congenital heart disease (CHD), portopulmonary disease and others), or induced by drugs and toxins (DTOX)^5^. Disease susceptibility includes genetic and environmental factors. Known risk genes underlie 70-80% of FPAH and ~10-40% of IPAH^6, 7^. However, the majority of non-familial cases remain genetically undefined.

Heterozygous germline mutations in bone morphogenetic protein receptor type 2 (*BMPR2*), a member of the transforming growth factor beta (TGF-β) superfamily, are the most common genetic cause of PAH ^8–10^. Similar frequencies of *BMPR2* mutations are observed across patient ethnicities and are present in 60-80% of familial cases ^11–14^. *BMPR2* mutations are observed in both child- and adult-onset PAH ^14^, and *BMPR2* mutation carriers exhibit a younger age-of-onset compared to non-carriers^7^. Mutations in the developmental transcription factor T-box 4 *(TBX4*) are more common in child-onset PAH, and *de novo* mutations in many different genes may explain ~19% of child-onset PAH ^14^. Germline mutations in other genes are individually rare causes of PAH. These include other genes in the TGF-β/BMP signaling pathway^15^, hereditary hemorrhagic telangiectasia (HHT) genes activin A receptor type II-like 1 (*ACVRL1*) and endoglin (*ENG*)^7^, eukaryotic initiation translation factor (*EIF2AK4*) associated with pulmonary veno-occlusive disease (PVOD)/pulmonary capillary hemangiomatosis (PCH)^16, 17^, caveolin-1 (*CAV1*)^18^, and channel genes including potassium two pore domain channel (*KCNK3*)^19^, ATP-binding cassette subfamily member 8 (*ABCC8*) ^20^ and voltage-dependent potassium channel 1.5 (*KCNA5*) ^21^.

New risk genes are emerging from large exome- and genome-wide sequencing studies. Rare mutations in SRY-related HMG-box transcription factor (*SOX17*), a key regulator of embryonic vasculogenesis, explain ~3.2% of APAH-CHD ^22^, and 0.7% of IPAH ^22, 23^. The UK NIHR BioResource – Rare Diseases PAH Study, utilizing ~1000 PAH cases of primarily adult-onset IPAH, identified an ATPase gene (*ATP13A3*), growth differentiation factor 2 (*GDF2*; also known as *BMP9*) and *SOX17* as risk genes contributing to 0.8-1.1% of cases ^23^. The low frequency of risk variants for each gene, except *BMPR2*, indicates that large numbers of individuals are required for further validation of rare risk genes and pathways, and to understand the natural history of each genetic subtype of PAH.

The National Biological Sample and Data Repository for PAH (AKA PAH Biobank) is a resource of biological specimens as well as clinical and genetic data generated for 2900 group 1 PAH patients to serve as a resource to the research community to enable larger-scale PAH studies. Herein, we performed targeted PAH gene and whole exome sequencing of 2572 cases from the PAH Biobank to identify and characterize frequencies and mutations in known PAH risk genes, identify new risk genes, and identify correlations between risk genes and clinical phenotypes.

## RESULTS

### Cohort Characteristics

Characteristics of the PAH Biobank cohort are shown in Table 1 and Supplementary Table 1. The cohort included 2572 cases: 43% IPAH, 48% APAH, 4% FPAH and 5% other PAH. The APAH cases included 722 associated with autoimmune CTDs (mostly scleroderma with few cases of rheumatoid arthritis, systemic lupus erythematosus and Sjogren’s syndrome), 268 with CHD, 139 with portopulmonary disease and 110 with other diseases (HHT, HIV and rare disorders). The “other PAH” group included 110 drug- and toxin-induced (DTOX) PAH, eleven non-familial PVOD/PCH cases and one persistent pulmonary hypertension of the newborn. The majority of cases (91.2%) were adult-onset with a cohort mean age-of-onset of 48 ±19 years (mean ± SD). However, there was an enrichment of child-onset cases (95/268, 37.2%, p<0.0001 by Chi-square) in the APAH-CHD subclass. As has been reported previously for adult populations^24^, there was an overall 3.7:1 ratio of females to males, with a 9:1 ratio for PAH associated with autoimmune disease and 1:1.2 ratio for the portopulmonary subclass. The genetic ancestries included European (72%), Hispanic (12%), African (11%), East Asian (2.7%) and South Asian (1.1%), fairly equally distributed amongst PAH subclasses. Within the APAH subclass, Africans were more likely to have disease associated with CTD (p=0.02) and less likely with CHD (p=0.0004) or portopulmonary disease (p=0.001), consistent with a previous report^25^. There was an enrichment of portopulmonary disease among patients of Hispanic ancestry (p=0.02).

**Table 1.** PAH Biobank cohort demographic and hemodynamic data.

|  | **ALL** | **IPAH** | **APAH** | **FPAH** | **Other\*** |
| --- | --- | --- | --- | --- | --- |
| **Total (%)** | 2572 | 1110 (43.2) | 1239 (48.2) | 101 (3.9) | 122 (4.7) |
| **Age-of-onset, n (%)** |  |  |  |  |  |
| Child (dx age <19) | 226 (8.8) | 94 (8.5) | 112 (9.0) | 15 (14.9) | 5 (4.1) |
| Adult (dx age ≥19) | 2345 (91.2) | 1015 (91.4) | 1127 (91.0) | 86 (85.1) | 117 (95.9) |
| Mean age | 48 ± 19 | 48 ± 18 | 49 ± 19 | 37 ± 15 | 47 ± 15 |
| **Gender, n (%)** |  |  |  |  |  |
| Female | 2023 (78.7) | 868 (78.2) | 996 (80.4) | 69 (68.3) | 90 (73.8) |
| Male | 549 (21.3) | 242 (21.8) | 243 (19.6) | 32 (31.7) | 32 (26.2) |
| Female:male ratio | 3.7:1 | 3.6:1 | 4.1:1 | 2.2:1 | 2.8:1 |
| **Ancestry, n (%)** |  |  |  |  |  |
| European | 1852 (72) | 809 (73.0) | 855 (69.0) | 89 (88.1) | 99 (81.2) |
| Hispanic | 315 (12.3) | 137 (12.3) | 156 (12.6) | 10 (9.9) | 12 (9.8) |
| African | 292 (11.4) | 117 (10.5) | 168 (13.6) | 1 (1) | 6 (4.9) |
| East Asian | 70 (2.7) | 25 (2.2) | 41 (3.3) | 0 | 4 (3.3) |
| South Asian | 28 (1.1) | 12 (1.1) | 15 (1.2) | 1 (1) | 0 |
| Others | 15 (0.58) | 10 (0.9) | 4 (0.3) | 0 | 1 (0.8) |
| **Hemodynamic parameters** |  |  |  |  |  |
| MPAP (mmHg) | 50 ± 14 | 52 ± 14 | 48 ± 14 | 58 ± 14 | 52 ± 13 |
| MPCW (mmHg) | 10 ± 4 | 10 ± 4 | 10 ± 4 | 10 ± 4 | 11 ± 4 |
| CO, Fick (L/min) | 4.5 ± 1.8 | 4.5 ± 1.7 | 4.6 ± 1.9 | 3.6 ± 1.0 | 4.2 ± 1.3 |
| PVR (Woods units) | 10.7 ± 7.0 | 11.2 ± 7.0 | 10.0 ± 7.1 | 14.9 ± 6.3 | 11.0 ± 6.6 |
| MAP (mmHg) | 90 ± 19 | 91 ± 20 | 90 ± 19 | 88 ± 16 | 94 ± 19 |
| MAP:MPAP | 1.9 ± 0.7 | 1.9 ± 0.7 | 2.0 ± 0.7 | 1.6 ± 0.5 | 1.9 ± 0.5 |

\*Other included 110 diet- and toxin-induced PAH, 11 non-familial pulmonary veno-occlusive disease/ pulmonary capillary hemangiomatosis and one persistent pulmonary hypertension of the newborn.
Abbreviations: MPAP, mean pulmonary arterial pressure; MPCW, mean pulmonary capillary wedge pressure; CO, cardiac output by Fick method; PVR, pulmonary vascular resistance; MAP, mean arterial pressure.

### Rare deleterious variants in established and recently-reported PAH risk genes

We screened for rare, predicted deleterious variants ((allele frequency <0.01% and likely gene damaging (LGD) or missense with REVEL score >0.5 (D-Mis), see Methods)) in eleven established PAH risk genes ^26–29^: *ACVRL1*, *BMPR1A*, *BMPR1B*, *BMPR2*, *CAV1*, *EIF2AK4*, *ENG*, *KCNK3*, *SMAD4*, *SMAD9*, and *TBX4* by targeted capture/sequencing, multiple ligation-dependent probe amplification (MLPA) (to evaluate deletions/duplications in *BMPR2*, *ACVRL1* and *ENG* only) and exome sequencing. We also screened the cohort for variants in seven recently-reported risk genes: *ABCC8*, *ATP13A3*, *GDF2/BMP9*, *KCNA5*, *KLF2*, *SMAD1*, and *SOX17*. Only fourteen percent of cases (n=349, 22% IPAH, 12% APAH, 55% FPAH, 11% other) carried rare predicted deleterious variants in these risk genes (Figure 1).

**Figure 1.**
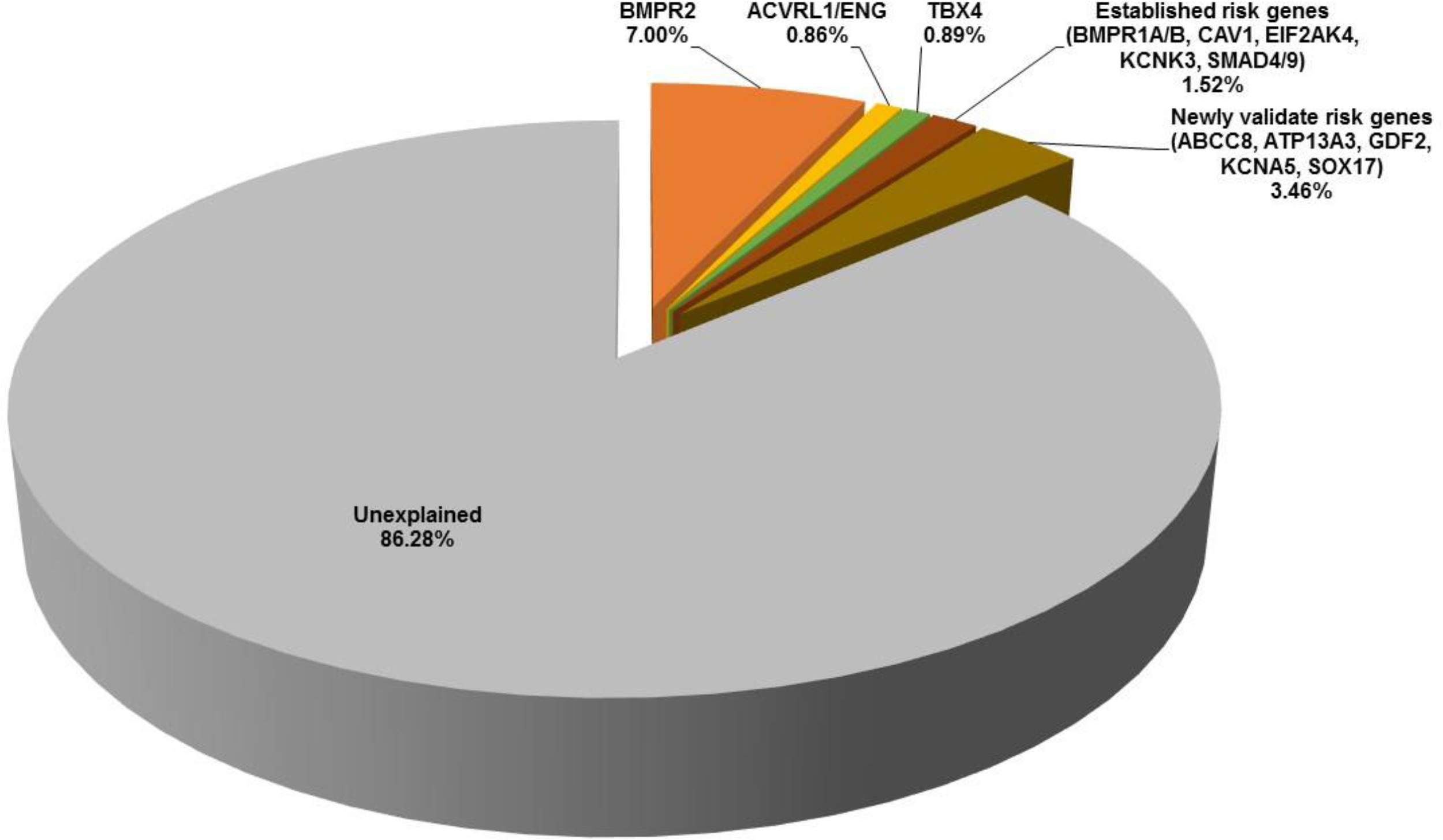
Contribution of known PAH risk genes in the PAH Biobank cohort (n=2572 cases). *BMPR2, ACVRL1/ENG, TBX4*; Other established risk genes included in the analysis: *BMPR1A*, *BMPR1B*, *CAV1*, *EIF2AK4*, *KCNK3*, *SMAD4* and *SMAD9;* Newly validated risk genes: *ABCC8*, *ATP13A1*, *GDF2*, *KCN5A*, *KLF2*, *SMAD1* and *SOX17*.

A complete list of cases carrying rare deleterious variants in established risk genes is provided in Supplementary Table 2. Not surprisingly, 68% of these cases carried variants in *BMPR2* (n=119 variants in 180 cases: 9% exon deletions, 65% LGD, 26% D-Mis). The age-of-onset for *BMPR2* variant carriers was 38 ± 15 years (mean ± SD), significantly younger than that of the whole cohort (p=1.1E-15, Mann-Whitney U test) but with a wide range of ages from 2 to 76 years (Figure 2). The second most common genetic cause was *TBX4*, accounting for approximately one percent of cases (n=23 cases with 22 variants: 12 LGD, 9 D-Mis and 1 in-frame deletion), the majority of whom (57%) had a diagnosis of IPAH. Although more than 90% of cases in the PAH Biobank cohort had adult-onset disease, only 48% of the *TBX4* variant carriers had adult-onset disease. The overall mean age-of-onset was 29 ± 25 years (Figure 2A), with a bimodal distribution and a significant enrichment of pediatric-onset cases compared to the whole PAH cohort (p=6.5E−08, RR=12.3, binomial test) (Figure 2B), consistent with previous findings^14^. Deleterious variants in nine additional genes were observed: *ACVRL1* (n=16 cases, including seven with HHT), *SMAD9* (13 cases), *CAV1* (10 cases), *ENG* (6 cases, including two with HHT), bi-allelic *EIF2AK4* (5 cases, including two with PVOD/PCH), *KCNK3* (3 cases), *BMPR1A* (4 cases), *SMAD4* (2 cases), and *BMPR1B* (2 cases). Four cases (2 IPAH, 2 FPAH) carried risk variants in *BMPR2* plus one other risk gene.

**Figure 2.**
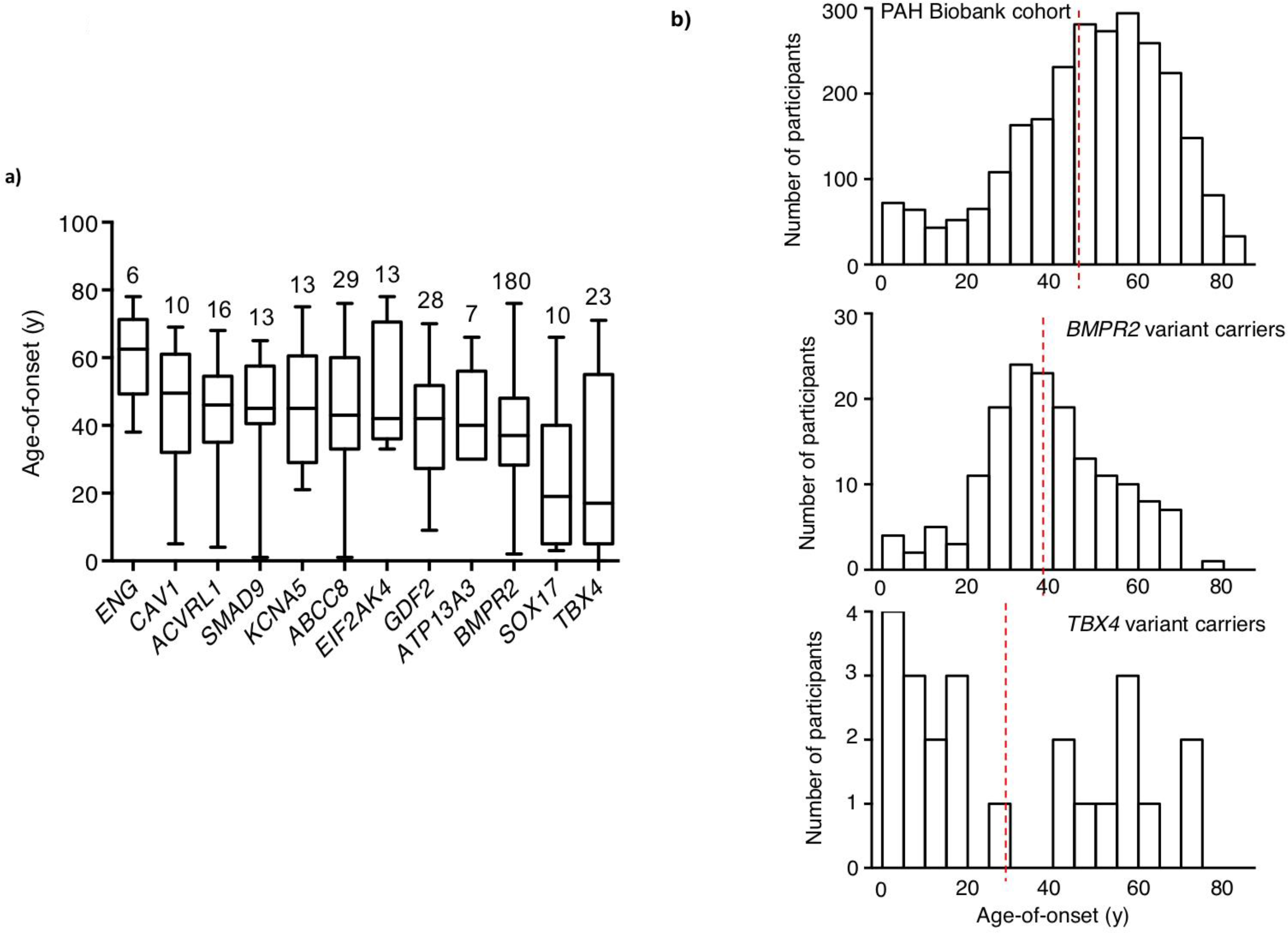
Age-of-disease onset for PAH Biobank cases with rare deleterious variants in known PAH risk genes. **a)** Box plots showing median, interquartile range and min/max values for age-of-disease onset (i.e. age at diagnostic right heart catheterization). The number of cases carrying variants for each gene is given above each box plot. Genes represented by less than four cases are not shown. **b)** Histogram plots showing age-of-onset distributions for the whole cohort (n=2572), *BMPR2* (n=180) or *TBX4* (n=23) variant carriers. Red vertical lines indicate the group means. *BMPR2* carriers had a younger mean age-of-onset (mean=37 y, SD=15; Mann-Whitney U test: p=1.1E-15) but no enrichment of child-onset cases (binomial test: p=1, RR=0.93) compared to the whole cohort, whereas *TBX4* carriers had a younger mean age-of-onset (mean=29 y, SD=25; Mann-Whitney U test: p=0.001) and significant enrichment of child-onset cases (binomial test: p=6.5E−08, RR=12.3) compared to the whole cohort.

A complete list of rare deleterious variants in newly-reported PAH risk genes is provided in Supplementary Table 3. Nearly two-thirds were variants in *ABCC8* (26 variants in 29 cases: all D-Mis) or *GDF2* (24 variants in 28 cases: 9 LGD, 15 D-Mis). The *ABCC8* variants occurred equally in IPAH and APAH cases (50:50) while the *GDF2* variants occurred primarily in IPAH cases (75%). Deleterious variants in the other new PAH risk genes were observed less frequently or not at all: *KCNA5* (n=13 cases), *SOX17* (10 cases), *ATP13A3* (7 cases), *SMAD1* (2 cases) and *KLF2* (0 cases). The mean age-of-onset for these risk gene variant carriers ranged from 41-46 years, with the exception of *SOX17* which had a mean age-of-onset of 26 years (Figure 2A), significantly younger than that of the whole cohort (p<0.003, Mann-Whitney U test). The female:male ratio among these patients was 4.2:1, similar to that of the whole cohort. Overall, 71% percent of the variants in known risk genes were novel.

Locations of the risk gene variants are shown in Supplementary Figures 1 and 2. For *BMPR2*, all but two of the D-Mis variants are located within the first 500 amino acids of the protein, mostly within the conserved activin and protein kinase domains (Supplementary Figure 1). While the LGD variants are also clustered within the activin and protein kinase domains, twenty-six variants carried by twenty-eight individuals are located downstream of these domains. For the other risk genes, the majority of D-Mis variants are also located in conserved protein domains (Supplementary Figure 2).

### Identification of novel PAH risk genes: KLK1 and GGCX

Our extensive genome screening efforts failed to identify rare deleterious variants in known risk genes for 86% of the PAH Biobank cases. To identify novel PAH risk genes, we performed a gene-based, case-control association analysis. To prevent confounding by genetic ancestry, we included only participants of European ancestry (cases: n=1832; controls: n= 7509 gnomAD WGS subjects and 5262 unaffected parents from the Pediatric Cardiac Genomics Consortium. To minimize technical batch effects of genotype data between cases and controls, we applied heuristic filters as described in Methods. We observed similar overall frequencies of rare synonymous variants in cases and controls (enrichment rate = 1.01, p-value=0.09), a class that is mostly neutral with respect to disease status (Supplemental Table 4). Further, a gene-level burden test confined to rare synonymous variants was consistent with a global null model (Supplementary Figure 3), indicating that technical batch effects would likely have minimal impact on genetic analyses. We then proceeded to test for gene-specific enrichment of rare deleterious variants (allele frequency <0.01%, LGD and D-Mis) in cases compared to controls. The use of in silico prediction tools to select deleterious missense variants can increase statistical power for rare variant association analyses^30^, but the optimal threshold for deleteriousness scores is often gene-specific^31^. To improve power, we implemented a rare variant burden test utilizing empirically-determined, gene-specific deleterious score thresholds, a “variable threshold test.” The association results across all protein-coding genes were generally consistent with the expectation under the null model (Figure 3). Across all PAH subclasses, only two genes exceeded the Bonferoni-corrected threshold for significance: *BMPR2* (p=1.0E−07, FDR=0.002) and *KLK1* (p=2.0E−07, FDR=0.002) (Figure 3). *KLK1* encodes kallikrein 1, also known as tissue kallikrein, involved in the regulation of systemic blood pressure and vascular remodeling but not previously associated with pulmonary hypertension ^32, 33^. Two other known risk genes, *ACVRL1* and *GDF2*, fell just below the cut-off for significance. We next repeated the analysis using 812 IPAH cases only (all European) and observed significant associations for *BMPR2* (p=1.0E−7, FDR=9.0E−04), *KLK1* (p=1.0E−7, FDR=9.0E−04), *GDF2* (p=3.0E−07, FDR=0.002) and *GGCX* (p=5.0E−07, FDR=0.002) (Figure 4). *GGCX* encodes gamma-glutamyl carboxylase, implicated in coagulation factor deficiencies and ectopic mineralization of soft tissues ^34^. These four genes were the only genes to reach genome-wide significance among IPAH cases. IPAH risk gene, *TBX4*, fell just below the cut-off for significance. All association results for the total cohort or IPAH alone, with p≤0.001, are listed in Figures 3 and 4, respectively. Analysis of the depth of sequencing coverage of the targeted regions in *KLK1* and *GGCX* indicated that nearly 100% of samples attained read depths of at least 15X, excluding the possibility that the associations were driven by coverage differences between cases and controls (Supplemental Figure 4).

**Figure 3.**
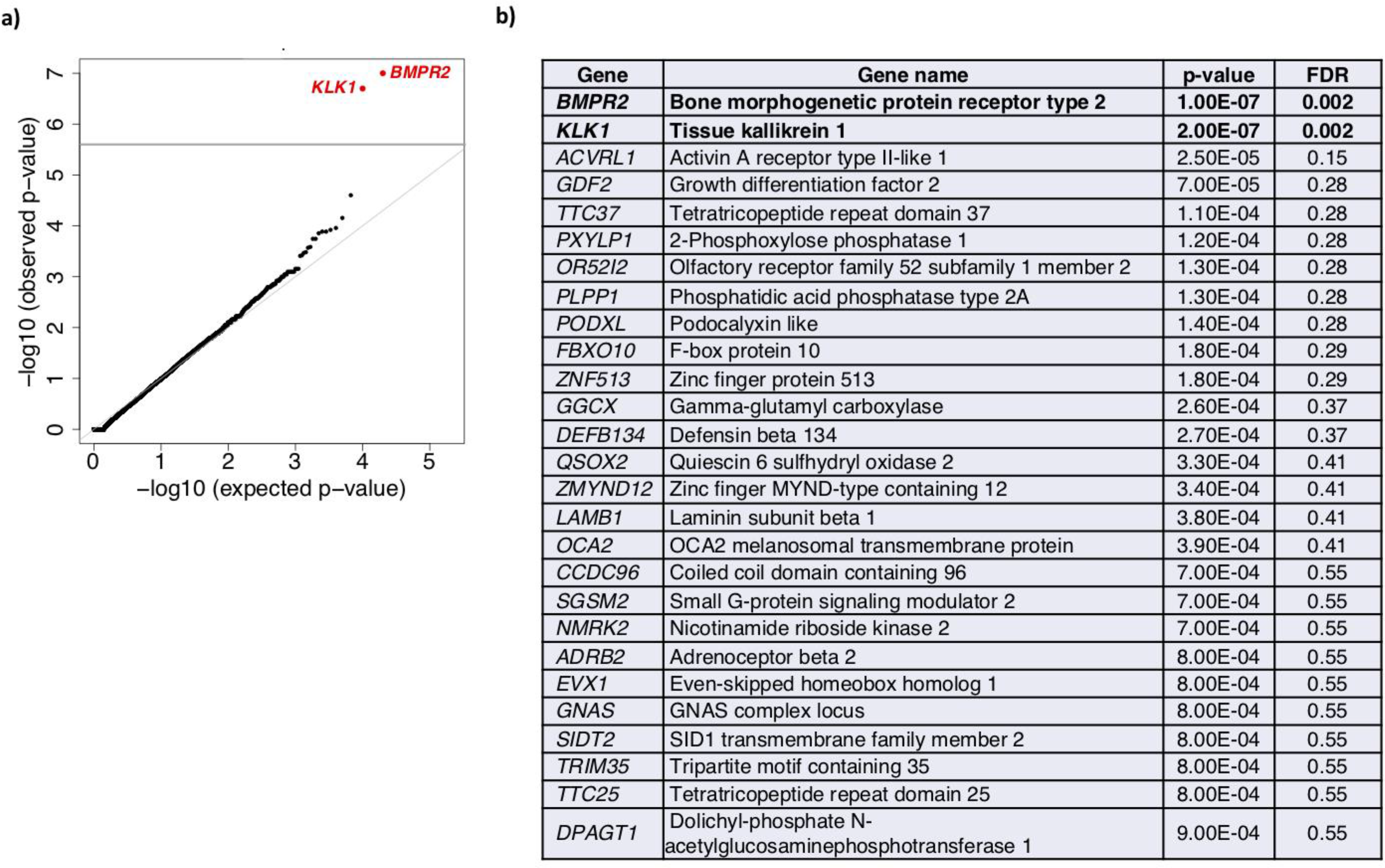
Gene-based association analysis using 1,832 European cases from all PAH subclasses and 12,771 European controls. **a)** Results of a binomial test confined to rare LGD and D-Mis (REVEL variable threshold) variants in 20,000 protein-coding genes. Horizontal gray line indicates the Bonferroni-corrected threshold for significance. **b)** Complete list of top association genes (p<=0.001).

**Figure 4.**
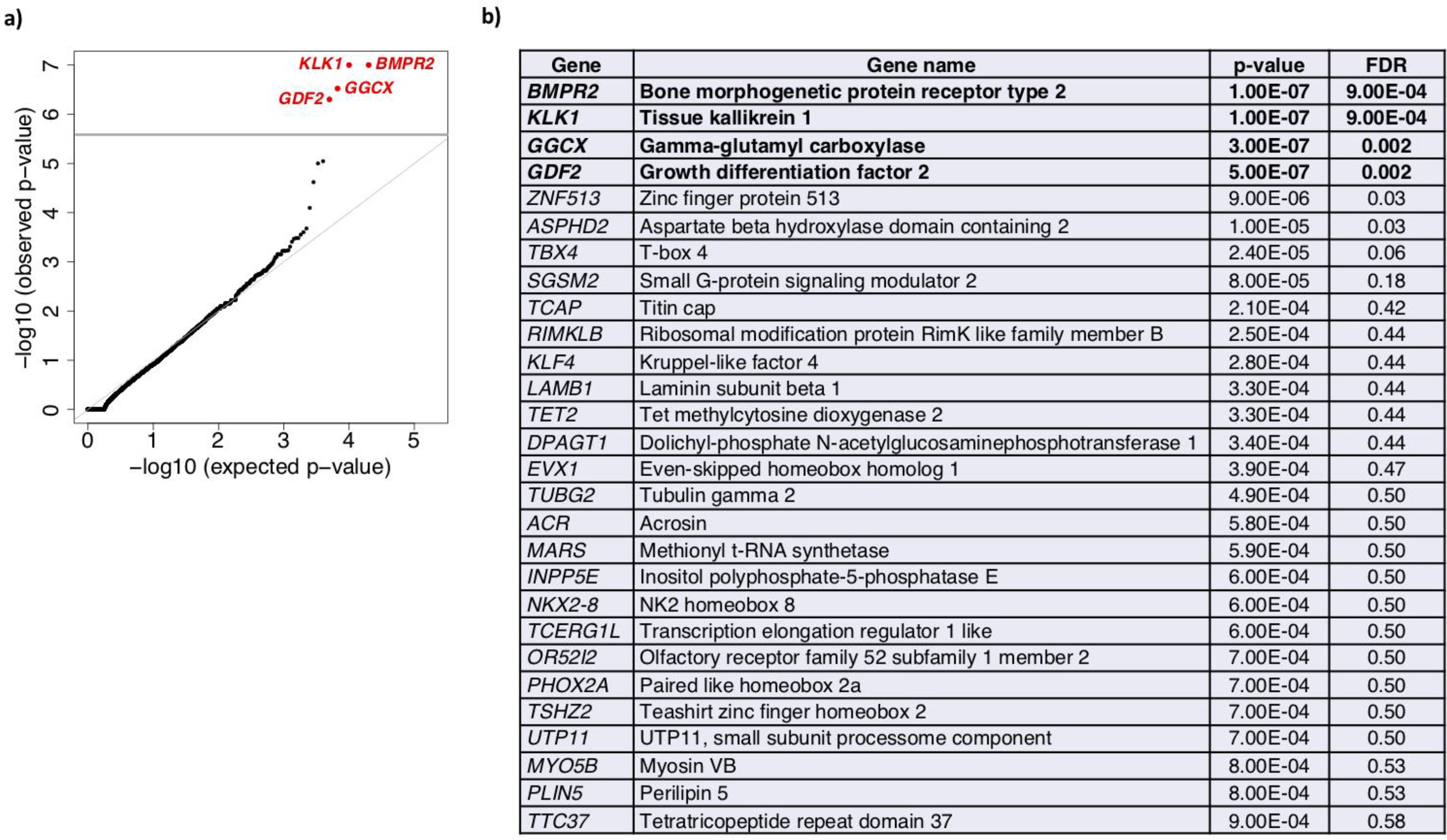
Gene-based association analysis using 812 European IPAH cases and 12,771 European controls. **a)** Results of a binomial test confined to rare LGD and D-Mis (REVEL variable threshold) variants in 20,000 protein-coding genes. Horizontal gray line indicates the Bonferroni-corrected threshold for significance. **b)** Complete list of top association genes (p<=0.001).

We next screened the entire PAH Biobank cohort, including participants of non-European ancestry, for rare deleterious variants in *KLK1* and *GGCX*. In total, 12 cases carried *KLK1* variants (10 IPAH, 2 APAH) and 28 cases carried *GGCX* variants (17 IPAH, 9 APAH, 1 FPAH, 1 unknown subclass) (Table 2). Most of the participants were of European ancestry, however, for *GGCX* there were also six cases of African and three cases of Hispanic ancestries. The mean age-of-onset was similar to that of the overall cohort for both genes (*KLK1*, 49 ± 6; *GGCX*, 49 ± 3). The variants for *KLK1* included four LGD (1 stop-gain, 2 frameshift, 1 splicing) and eight D-Mis; variants for *GGCX* included six LGD (5 stop-gain, 1 frameshift), twenty-one D-Mis and one in-frame deletion. Three *KLK1* (1 LGD, 2 D-Mis) and five *GGCX* (1 LGD, 4 D-Mis) variants were recurrent in the cohort. Locations of the variant amino acid residues are shown in Figure 5. All but one of the *KLK1* and two of the *GGCX* missense variants, as well as the in-frame deletion, occur in conserved enzymatic domains.

**Table 2.** Rare, predicted deleterious *KLK1* and *GGCX* variants* among 2,572 PAH cases. Participants were heterozygous for the indicated variants.

| Participant ID | Gender | Age at dx (y) | PAH subclass | Ancestry | Gene** | Nucleotide change | Amino acid change | Variant type | MAF (ExAC) | CADD score | Revel score |
| --- | --- | --- | --- | --- | --- | --- | --- | --- | --- | --- | --- |
| 08-022 | F | 60 | IPAH | EUR | *KLK1* | c.46+1G>T | p.(=) | splicing | . | 24 | . |
| 10-096 | F | 68 | IPAH | EUR | *KLK1* | c.60dup | p.Ile21Aspfs*12 | frameshift | 4.29E-05 | . | . |
| 28-049 | F | 36 | APAH-CHD | EUR | *KLK1* | c.60dup | p.Ile21Aspfs*12 | frameshift | 4.29E-05 | . | . |
| 06-058 | M | 13 | IPAH | EUR | *KLK1* | c.70C>T | p.Arg24Trp | D-Mis | 8.47E-06 | 26 | 0.56 |
| 13-002 | F | 71 | IPAH | EUR | *KLK1* | c.70C>T | p.Arg24Trp | D-Mis | 8.47E-06 | 26 | 0.56 |
| 06-007 | M | 26 | IPAH | EUR | *KLK1* | c.113G>A | p.Trp38* | stopgain | 8.30E-06 | 35 | . |
| 12-061 | F | 51 | IPAH | EUR | *KLK1* | c.469G>A | p.Gly157Ser | D-Mis | 9.36E-05 | 29 | 0.72 |
| 14-018 | M | 61 | IPAH | EUR | *KLK1* | c.469G>A | p.Gly157Ser | D-Mis | 9.36E-05 | 29 | 0.72 |
| 17-075 | F | 82 | APAH-CTD | EUR | *KLK1* | c.469G>A | p.Gly157Ser | D-Mis | 9.36E-05 | 29 | 0.72 |
| 18-026 | F | 37 | IPAH | EUR | *KLK1* | c.644G>A | p.Gly215Glu | D-Mis | . | 30 | 0.85 |
| 19-013 | F | 51 | IPAH | EUR | *KLK1* | c.650C>T | p.Pro217Leu | D-Mis | 2.52E-05 | 29 | 0.60 |
| 19-033 | F | 37 | IPAH | EUR | *KLK1* | c.689G>C | p.Trp230Ser | D-Mis | 8.26E-06 | 25 | 0.50 |
| 06-014 | M | 35 | FPAH | EUR | *GGCX* | c.137C>G | p.Ser46Cys | D-Mis | 5.77E-05 | 23 | 0.70 |
| 04-020 | F | 36 | IPAH | EUR | *GGCX* | c.G203G>C | p.Arg68Pro | D-Mis | . | 35 | 0.96 |
| 12-207 | F | 43 | IPAH | EUR | *GGCX* | c.G203G>C | p.Arg68Pro | D-Mis | . | 35 | 0.96 |
| 32-003 | M | 81 | IPAH | EUR | *GGCX* | c.248G>A | p.Arg83Gln | D-Mis | . | 34 | 0.92 |
| 32-008 | F | 36 | IPAH | AFR | *GGCX* | c.322C>T | p.Arg108Cys | D-Mis | 1.65E-05 | 31 | 0.55 |
| 08-013 | F | 66 | APAH-CTD | EUR | *GGCX* | c.646_647delinsCA | p.Val216Gln | in-frame | . | 31 | *** |
| 26-036 | F | 52 | IPAH | EUR | *GGCX* | c.722T>C | p.Phe241Ser | D-Mis | . | 33 | 0.94 |
| 30-031 | F | 55 | IPAH | EUR | *GGCX* | c.722T>C | p.Phe241Ser | D-Mis | . | 33 | 0.94 |
| 04-029 | F | 60 | IPAH | EUR | *GGCX* | c.734T>A | p.Leu245* | stopgain | . | 40 | . |
| 04-087 | F | 54 | IPAH | EUR | *GGCX* | c.763G>A | p.Val255Met | D-Mis | 1.65E-05 | 34 | 0.86 |
| 34-005 | M | 66 | IPAH | EUR | *GGCX* | c.763G>A | p.Val255Met | D-Mis | 1.65E-05 | 34 | 0.86 |
| 11-004 | F | 24 | IPAH | HIS | *GGCX* | c.763G>A | p.Val255Met | D-Mis | 1.65E-05 | 34 | 0.86 |
| 22-108 | F | 40 | APAH-HIV | AFR | *GGCX* | c.899C>T | p.Ser300Phe | D-Mis | 2.53E-05 | 28 | 0.82 |
| 28-110 | F | 56 | IPAH | AFR | *GGCX* | c.899C>T | p.Ser300Phe | D-Mis | 2.53E-05 | 28 | 0.82 |
| 28-096 | F | 23 | IPAH | EUR | *GGCX* | c.938_939del | p.Pro313Argfs*33 | frameshift | 1.00E-04 | . | . |
| 08-046 | F | 53 | APAH-Porto | EUR | *GGCX* | c.950G>A | p.Arg317Gln | D-Mis | 1.67E-05 | 33 | 0.81 |
| 12-205 | F | 55 | IPAH | EUR | GGCX | c.1017_1018insT | p.Ser340* | stopgain | . | . | . |
| 15-008 | F | 14 | APAH-CHD | EUR | GGCX | c.1075C>T | p.Arg359Cys | D-Mis | . | 28 | 0.76 |
| 21-037 | F | 45 | APAH-CTD | AFR | GGCX | c.1128C>G | p.Phe376Leu | D-Mis | . | 27 | 0.85 |
| 06-039 | F | 28 | IPAH | EUR | GGCX | c.1224C>A | p.His408Gln | D-Mis | 8.24E-06 | 23 | 0.75 |
| 37-004 | F | 48 | IPAH | EUR | GGCX | c.1249G>A | p.Asp417Asn | D-Mis | 1.65E-05 | 26 | 0.72 |
| 30-034 | F | 49 | APAH-CTD | HIS | GGCX | c.1304G>A | p.Arg435Gln | D-Mis | 8.24E-06 | 29 | 0.67 |
| 14-029 | M | 48 | IPAH | EUR | GGCX | c.1306C>T | p.Arg436* | stopgain | 3.30E-05 | 41 | . |
| 37-010 | F | 77 | APAH-CTD | EUR | GGCX | c.1306C>T | p.Arg436* | stopgain | 3.30E-05 | 41 | . |
| 11-090 | F | 47 | APAH | AFR | GGCX | c.1465G>A | p.Val489Met | D-Mis | 2.47E-05 | 26 | 0.68 |
| 05-013 | M | 63 | APAH-Porto | HIS | GGCX | c.1480T>G | p.Ser494Ala | D-Mis | . | 26 | 0.84 |
| 28-033 | F | 51 | IPAH | EUR | GGCX | c.1758C>G | p.Tyr586* | stopgain | . | 45 | . |
| 17-033 | F | 74 | APAH-CTD | AFR | GGCX | c.1772C>T | p.Thr591Met | D-Mis | 3.30E-05 | 29 | 0.83 |

\*Rare, deleterious variants defined as gnomAD AF $\leq 1.00E-04$ and REVEL $>0.5$ .
\*\**KLK1* transcript NM\_002257.3 and *GGCX* transcript NM\_000821.6.
\*\*\*REVEL score could not be computed for this 2-nt substitution because machine learning is based on 1-nt substitutions. Inclusion in the table was based on REVEL $>0.9$ for single nt substitution and PROVEAN=deleterious for 2-nt substitution.

**Figure 5.**
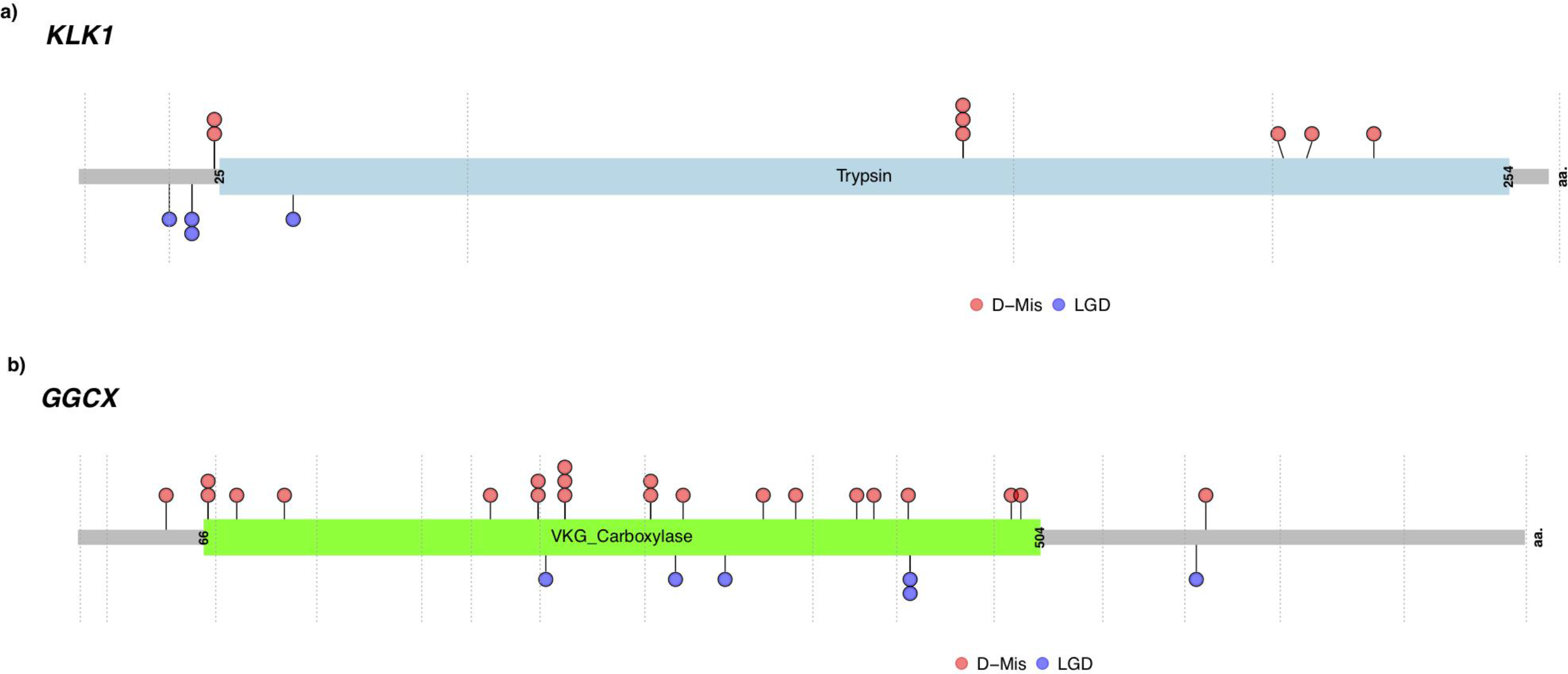
Locations of rare, predicted deleterious variants in *KLK1* (a) and *GGCX* (b) across the PAH Biobank cohort (n=2572 cases). Locations are provided within the two-dimensional protein structures. The numbers of variants at each amino acid position are indicated along the y-axes. The vertical gray lines indicate exon borders. D-MIS, predicted damaging missense; LGD, likely-gene-disrupting (stopgain, frameshift, splicing).

*KLK1* belongs to a contiguous gene family cluster on chromosome 19 encoding fifteen distinct peptidases. While some have highly restricted expression patterns (ie *KLK2* and *KLK3* in prostate), others are widely expressed^35^. Nine of the family members, including *KLK1*, are expressed in lung and have been implicated in various lung diseases – inflammatory respiratory diseases, viral infections, and cancers^36^. We tested for enrichment of rare deleterious variants in the gene-set expressed in lung and observed a significant enrichment of LGD+D-Mis variants in European cases compared to controls (enrichment rate=2.1, p=0.004) (Supplementary Table 5A). We then performed gene-specific association analyses to determine which genes were contributing to the enrichment. The associations were stronger for IPAH than all PAH; while *KLK1* was the only gene to exceed the Bonferroni-corrected threshold for significance (OR=13.9, p=2.00E−07 for all PAH; OR=26.2, p=1.00E−07 for IPAH), five additional family members had an enrichment rate of rare deleterious variants greater than 2.0 for IPAH (Supplementary Table 5B).

### Clinical phenotypes of KLK1 and GGCX variant carriers

Hemodynamic measurements at the time of PAH diagnosis for individual carriers of *KLK1* and *GGCX* variants are provided in Table 3. Clinical phenotypes of IPAH participants with *KLK1* or *GGCX* variants did not differ from that of other IPAH cases without variants in known risk genes (Supplementary Table 6). Overall, participants with predicted deleterious variants in either gene exhibited less severe clinical phenotypes compared to participants with variants in *BMPR2*. Carriers of both *KLK1* and *GGCX* variants were older at PAH onset and had decreased mean pulmonary arterial pressure, increased cardiac output and decreased pulmonary vascular resistance compared to *BMPR2* carriers (Table 3). Furthermore, both *KLK1* and *GGCX* carriers had increased ratios of mean (systemic) arterial pressure to mean pulmonary artery pressure compared to *BMPR2* carriers (MAP:MPAP, Table 3).

**Table 3.** Clinical phenotypes of *KLK1* and *GGCX* variant carriers at PAH diagnosis, and compared to mean phenotypes of *BMPR2* variant carriers.

| Participant ID | Gender | PAH subclass | Gene | Age at dx (y) | MPAP (mmHg) | MPCW (mmHg) | CO, Fick (L/min) | PVR (Woods units) | MAP (mmHg) | MAP: MPAP |
| --- | --- | --- | --- | --- | --- | --- | --- | --- | --- | --- |
| 08-022 | F | IPAH | *KLK1* | 60 | 55 | 7 | 2.6 | 18.46 | 98 | 1.78 |
| 10-096 | F | IPAH | *KLK1* | 68 | 46 | 9 | 3.95 | 9.37 | NA | NA |
| 28-049 | F | APAH-CHD | *KLK1* | 36 | 61 | 5 | 3.77 | 14.85 | 82 | 1.34 |
| 06-058 | M | IPAH | *KLK1* | 13 | 38 | 8 | 8.2 | 3.66 | 92 | 2.42 |
| 13-002 | F | IPAH | *KLK1* | 71 | 44 | 9 | 3.7 | 9.46 | NA | NA |
| 06-007 | M | IPAH | *KLK1* | 26 | 41 | 11 | 5.8 | 5.17 | 97 | 2.37 |
| 12-061 | F | IPAH | *KLK1* | 51 | 53 | 13 | NA | NA | 106 | 2.00 |
| 14-018 | M | IPAH | *KLK1* | 61 | 37 | 6 | 4.38 | 7.08 | 98 | 2.65 |
| 17-075 | F | APAH-CTD | *KLK1* | 82 | 27 | 7 | 5.23 | 3.82 | NA | NA |
| 18-026 | F | IPAH | *KLK1* | 37 | 73 | 15 | NA | NA | NA | NA |
| 19-013 | F | IPAH | *KLK1* | 51 | 42 | 13 | 3.6 | 8.06 | 112 | 2.67 |
| 19-033 | F | IPAH | *KLK1* | 37 | 34 | 8 | 5.53 | 4.70 | 93 | 2.74 |
| Mean ± SD, *KLK1* | | | | 49 ± 20 | 46 ± 13 | 9 ± 3 | 4.7 ± 1.6 | 8.5 ± 4.9 | 97 ± 9 | 2.3 ± 0.5 |
| n, *KLK1* |  |  |  | 12 | 12 | 12 | 10 | 10 | 8 | 8 |
| Mean ± SD, *BMPR2* | | | | 38 ± 15 | 59 ± 12 | 10 ± 4 | 3.7 ± 1.3 | 15.3 ± 7.3 | 90 ± 17 | 1.6 ± 0.4 |
| n, *BMPR2* |  |  |  | 181 | 175 | 172 | 123 | 120 | 114 | 114 |
| p-value, *KLK1* vs *BMPR2* |  |  |  | 0.014 | 0.0007 | NS | 0.02 | 0.004 | NS | <0.0001 |
| 06-014 | M | FPAH | *GGCX* | 35 | 56 | 7 | 2.2 | 22.27 | 103 | 1.84 |
| 04-020 | F | IPAH | *GGCX* | 36 | 78 | 8 | 2.6 | 26.92 | 71 | 0.91 |
| 12-207 | F | IPAH | *GGCX* | 43 | 68 | 7 | 2.63 | 23.19 | NA | NA |
| 32-003 | M | IPAH | *GGCX* | 81 | 31 | 4 | 4.51 | 5.99 | NA | NA |
| 32-008 | F | IPAH | *GGCX* | 36 | 49 | 15 | 6.13 | 5.55 | 87 | 1.78 |
| 08-013 | F | APAH-CTD | *GGCX* | 66 | 40 | 9 | 5.8 | 5.34 | 120 | 3.00 |
| 26-036 | F | IPAH | *GGCX* | 52 | 70 | 9 | NA | NA | 86 | 1.23 |
| 30-031 | F | IPAH | *GGCX* | 55 | 56 | 15 | NA | NA | 121 | 2.16 |
| 04-029 | F | IPAH | *GGCX* | 60 | 51 | 8 | 5.53 | 7.78 | NA | NA |
| 04-087 | F | IPAH | *GGCX* | 54 | 40 | 14 | 5.64 | 4.61 | 115 | 2.88 |
| 34-005 | M | IPAH | GGCX | 66 | 43 | 14 | 6.85 | 4.23 | 83 | 1.93 |
| 11-004 | F | IPAH | GGCX | 24 | 77 | 11 | NA | NA | NA | NA |
| 22-108 | F | APAH-HIV | GGCX | 40 | 78 | 13 | 5.7 | 11.40 | 94 | 1.21 |
| 28-110 | F | IPAH | GGCX | 56 | 78 | NA | 3.6 | NA | 85 | 1.09 |
| 28-096 | F | IPAH | GGCX | 23 | 65 | 7 | 4.8 | 12.08 | NA | NA |
| 08-046 | F | APAH-Porto | GGCX | 53 | 44 | 10 | 6.23 | 5.46 | 88 | 2.00 |
| 12-205 | F | IPAH | GGCX | 55 | 52 | 8 | NA | NA | NA | NA |
| 15-008 | F | APAH-CHD | GGCX | 14 | 60 | 12 | 2.5 | 19.20 | 62 | 1.03 |
| 21-037 | F | APAH-CTD | GGCX | 45 | 31 | 12 | 5.26 | 3.61 | NA | NA |
| 06-039 | F | IPAH | GGCX | 28 | 48 | 11 | 4.6 | 8.04 | 81 | 1.69 |
| 37-004 | F | IPAH | GGCX | 48 | 49 | 18 | 3.6 | 8.61 | 67 | 1.37 |
| 30-034 | F | APAH-CTD | GGCX | 49 | 38 | 17 | NA | NA | NA | NA |
| 14-029 | M | IPAH | GGCX | 48 | 50 | 14 | NA | NA | NA | NA |
| 37-010 | F | APAH-CTD | GGCX | 77 | 28 | 6 | NA | NA | 74 | 2.64 |
| 11-090 | F | APAH | GGCX | 47 | NA | NA | NA | NA | NA | NA |
| 05-013 | M | APAH-Porto | GGCX | 63 | 33 | 6 | 6.31 | 4.28 | 119 | 3.61 |
| 28-033 | F | IPAH | GGCX | 51 | 61 | NA | 3.6 | NA | 85 | 1.39 |
| 17-033 | F | APAH-CTD | GGCX | 74 | 45 | 7 | 3.69 | 10.30 | 88 | 1.96 |
| Mean ± SD, *GGCX* | | | | 49 ± 16 | 53 ± 15 | 10 ± 4 | 4.6 ± 1.4 | 10.5 ± 7.4 | 91 ± 18 | 1.9 ± 0.8 |
| n, *GGCX* |  |  |  | 28 | 27 | 25 | 20 | 18 | 18 | 18 |
| Mean ± SD, *BMPR2* | | | | 38 ± 15 | 59 ± 12 | 10 ± 4 | 3.7 ± 1.3 | 15.3 ± 7.3 | 90 ± 17 | 1.6 ± 0.4 |
| n, *BMPR2* |  |  |  | 181 | 175 | 172 | 123 | 120 | 114 | 114 |
| p-value, *GGCX* vs *BMPR2* |  |  |  | <0.0001 | 0.02 | NS | 0.004 | 0.01 | NS | 0.007 |

Abbreviations: Age at dx, participant age at diagnosis/right heart catheterization; MPAP, mean pulmonary arterial pressure; MPCW, mean pulmonary capillary pressure; CO, cardiac output by the Fick method; PVR, pulmonary vascular resistance; MAP, mean arterial pressure.

A known *KLK1* single nucleotide polymorphism conferring at least partial loss of function occurs with high frequency in the general population ^37, 38^. The c.230G>A;p.R77H SNP (formerly called c.230G>A;p.R53H) has been associated with decreased urinary kallikrein activity and aberrant flow-mediated arterial remodeling but not systemic hypertension ^37, 38^. We screened the PAH Biobank cohort for the c.230G>A;p.R77H SNP and compared the cohort allele frequency with the frequency observed in gnomAD. No enrichment was observed in the PAH Biobank cohort (Supplementary Table 7), and none of the carriers of rare deleterious *KLK1* variants also carried the c.230G>A;p.R77H SNP. Thus, the observed association of rare, deleterious *KLK1* variants with PAH and associated phenotypes could not be explained by coincident occurrence of the common SNP.

## DISCUSSION

Using exome sequencing of a large PAH Biobank cohort recruited by 28 participating centers, followed by rare deleterious variant identification and gene-based association analysis, we identified *KLK1* and *GGCX* as novel candidate genes for PAH. These candidate risk genes suggest new pathogenic mechanisms outside of the TGF-β/BMPR2 signaling pathway. We showed that carriers of rare, predicted deleterious variants in *KLK1* or *GGCX* have less severe clinical phenotypes compared to carriers of *BMPR2* variants. In addition, we identified 252 novel rare deleterious variants in seventeen known PAH risk genes and confirmed the importance of *TBX4* and *SOX17* in early-onset disease as well as the genome-wide association of *GDF2* with IPAH.

The results of our study replicate some, but not all, of the recently-reported findings from the UK NIHR BioResource – Rare Diseases PAH Study. Similar to that cohort ^23^, *GDF2* reached significant genome-wide association among 812 IPAH cases of European ancestry in the PAH Biobank; with additional variants observed in a small number of APAH and FPAH participants. In total, we identified twenty-four variants, only two of which had been reported previously. *GDF2* encodes a well-characterized ligand for BMPR2, and these data further confirm an important role for *GDF2* in IPAH, as well as other PAH subclasses. Similar to the UK cohort, as well as our previous report of a cohort enriched in APAH-CHD cases^22^, we observed a low frequency of *SOX17* variants (0.4%) in the PAH Biobank likely due, at least in part, to the paucity of APAH-CHD cases in both Biobank cohorts. Interestingly, a genome-wide association study of common SNPs involving both the PAH Biobank and the UK NIHR BioResource – Rare Diseases PAH Study identified SNPs in a putative endothelial-acting enhancer region of *SOX17* in PAH^39^, suggesting that common variants may play an important role in susceptibility to PAH. Neither *ATP13A3* nor *AQP1* reached genome-wide significance in our study. We analyzed *ATP13A1* rare deleterious variants and identified seven cases with novel variants. *AQ1* not only failed to reach genome-wide significance but also was not among the expanded list of genes with p≤0.001 for either the whole PAH cohort or IPAH alone. Based on the small relative risks and associated confidence intervals from the UK NIHR BioResource – Rare Diseases PAH Study (RR=0.37, CI: 0.06-1.47 for *ATP13A3*; RR=0.19, CI: 0.004-1.53 for *AQP1*), it was not unexpected that that by chance we would observe no association with these genes and PAH.

The new PAH candidate risk genes identified in the current study, *KLK1* and *GGCX*, are both expressed in lung and vascular tissues, play important roles in vascular hemodynamics and inflammation, but have not been implicated in PAH previously. *KLK1*, also known as tissue kallikrein 1, is a major component of the kallikrein-kinin system that, together with the renin-angiotensin system, regulates blood pressure and cardiovascular function. In rodent and *in vitro* studies, *KLK1* is constitutively expressed by endothelial cells, and endothelial activation leads to release of active protease, matrix degradation, smooth muscle cell migration and vascular sprouting^32^, processes relevant to PAH. Gene delivery of tissue *KLK1* via adenoviral vectors, protein infusion or genetically-modified stem cells has shown beneficial effects in multiple models of vascular diseases^40^. *KLK1* is part of a highly-conserved, serine protease subfamily. We observed enrichment of rare deleterious variants in a gene-set of nine *KLK* genes expressed in lung, suggesting candidate genes for further investigation including *KLK12* which may play a role in angiogenesis via indirect regulation of vascular endothelial growth factor ^41^. The potential for gene delivery of *KLKs* as a treatment for PAH should prioritize this gene family for functional studies.

*GGCX* encodes gamma glutamyl carboxylase, responsible for the post-translational modification of vitamin-K dependent proteins involved in coagulation, soft tissue mineralization, inflammation, bone formation and cell proliferation ^34^. Homozygous mutations in *GGCX* cause vitamin K-dependent clotting factor deficiency (MIM #277450) as well as pseudoxanthoma elasticum (MIM #264800), an ectopic mineralization disorder. None of the PAH Biobank *GGCX* heterozygous variant carriers had diagnoses of bleeding disorders or pseudoxanthoma elasticum. *Ggcx*^*−/−*^ mice die pre- or perinatally due to massive bleeding but heterozygotes are viable^42^. Thus, the heterozygous knock-out mouse may provide a model for testing the effect of *Ggcx* on PAH phenotypes.

The differences in etiology, clinical course and prognosis for child- vs adult-onset PAH is an area of active investigation. Previous studies have implicated *BMPR2*, *TBX4* and *SOX17* in child-onset disease. In our large PAH Biobank cohort, *BMPR2* variant carriers exhibited a shift towards younger age-of onset, but the overall age distribution was similar to that of the whole cohort. *TBX4* exhibited a bi-modal distribution with significant enrichment of variants among pediatric-onset cases. Consistent with our previous report of *SOX17* in APAH-CHD, *SOX17* carriers in the PAH Biobank also had a relatively young mean age-of-onset (26 y). The hypothesis that pediatric PAH is linked to lung growth and development^43^ is consistent with roles for *TBX4* and *SOX17*, prominent developmental transcription factors^44^, in early-onset disease.

In summary, we have identified *KLK1* and *GGCX* as new candidate risk genes for PAH, accounting for ~0.4% and 0.9% of PAH Biobank cases, respectively. These genes require further genetic validation and functional assessment with the goal of providing new therapeutics for this lethal vasculopathy. The growing list of PAH risk genes and variants indicate that exome sequencing may be useful in families with PAH if no genetic cause is identified with panel gene testing. Furthermore, genomic studies of larger international consortia will be necessary to better clinically characterize these rare genetic subtypes of PAH.

## METHODS

### Participants

The PAH Biobank is housed and maintained at the Cincinnati Children’s Hospital Medical Center (CCHMC). Thirty-eight North American PH Centers participate in the PAH Biobank to identify and enroll patients meeting eligibility criteria. Each enrolling center also completes an electronic case report form with clinical data for each patient enrolled. Participants are diagnosed according to the World Health Organization PH group I classification^5^ and the diagnosis of PAH is confirmed by medical record review including right heart catheterization. The cohort for this genetic analysis included 2534 singletons, 19 duos (proband and one unaffected parent) and 19 trios (proband and two unaffected biological parents). Written informed consent (and assent when appropriate) was obtained from participants or parents/legal guardians under a protocol approved by the institutional review board at CCHMC as well as those at each of the participating PH Centers. Written informed consent for publication was obtained at enrollment. The data and resources are made available to the research community for hypothesis-driven projects via an application process (www.pahbiobank.org). A subset including 183 affected participants were included in previous publications from our group^14, 20, 22^.

### Targeted sequencing and multiplex ligation-dependent probe amplification (MLPA)

After proper informed consent, blood samples were collected and shipped to CCHMC for processing and generation of genetic data including panel sequencing of up to twelve genes, SNP genotyping using the Illumina OMNI5-4 Beadchip and limited MLPA dosage data. Targeted Next-Generation sequencing was performed with 250 ng DNA using the Illumina Tru-seq Custom Amplicon system (Illumina, USA) according to manufacturer’s instructions. Custom amplicons were designed with Illumina’s DesignStudio for the coding sequence of *BMPR2*, *ACVRL1*, *ENG*, *CAV1*, *SMAD9*, *KCNK3*, and *EIF2AK4*, for which all participants were sequenced. *ABCC8*, *GDF2*, *KCNA5*, *SMAD4*, and *TBX4* were added to the panel later and a subset of 739 were also sequenced for these genes. Each sample was sequenced using the Illumina MiSeq^®^ instrument with paired-end 250 nucleotide read lengths. Demultiplexing, base calling and alignment were executed using the default Illumina TruSeq Amplicon Workflow. Fastq files were aligned and visualized with NextGENe (SoftGenetics, USA). Variants were confirmed via Sanger sequencing on an ABI 3730xl DNA analyzer (Applied Biosystems, USA).

MLPA was performed with 100 ng of genomic DNA according to manufacturer’s instructions using the P093 Salsa MLPA probe sets (MRC-Holland, Amsterdam, The Netherlands). This probe set includes probes for all exons of *BMPR2*, *ALK1*, and *ENG*. Probe amplification products were run on an ABI 3730xl DNA Analyzer using GS500 size standard (Applied Biosystems). MLPA peak data was imported into Coffalyser (MRC-Holland) for quality checks and dosage ratio analysis. A dosage ratio value of ≤0.7 was used as the boundary for deletions, and ≥1.35 was used as the boundary for duplications.

### Whole exome sequencing (WES)

Exome sequencing was performed for the entire cohort in collaboration with the Regeneron Genetics Center (RGC). In brief, genomic DNA was prepared with a customized reagent kit from Kapa Biosystems and captured using Integrated DNA Technologies xGen lockdown probes. All samples were sequenced on the Illumina HiSeq 2500 platform using v4 chemistry, generating 76 bp paired-end reads. At least 90% of targeted regions have read depth coverage ≥15x for all exome sequencing samples.

### WES Data analysis

We used a previously established bioinformatics procedure ^45^ to process and analyze exome sequence data. Specifically, we used BWA-MEM (Burrows-Wheeler Aligner)^46^ to map and align paired-end reads to the human reference genome (version GRCh38/hg18), Picard MarkDuplicates to identify and flag PCR duplicate reads and GATK HaplotypeCaller (version 3.5)^47, 48^ to call genetic variants. We used heuristic filters to minimize technical artifacts, excluding variants that met any of the following criteria: missingness >10%, minimum read depth ≤8 reads, allele balance ≤25%^49^, and genotype quality <90. We obtained gnomAD whole genome sequence (WGS) as part of the control set. Only variants with FILTER “PASS” in gnomAD WGS (data release v 2.02) and restricted to the xGen captured protein coding region were included in the analysis. We used ANNOVAR^50^ to annotate the variants and aggregate information about allele frequencies (AF) and *in silico* predictions of deleteriousness. Rare variants were defined as AF ≤0.01% in both ExAC and gnomAD WES datasets. An exception was made for recessive inheritance of *EIF2AK4* variants, in which the AF cut-off was ≤1% Deleterious variants were defined as likely-gene-disrupting (including premature stop-gain, frameshift indels, canonical splicing variants and exon deletions) or predicted damaging missense with REVEL score >0.5 (D-Mis), as previously described ^22, 51^. Insertion/deletion variants were manually inspected using Integrative Genome Viewer (IGV). All variants in known PAH risk genes, included in Supplementary Tables 2 and 3, are deposited in ClinVar (pending).

### Statistical analysis

To identify novel candidate risk genes, we performed a gene-based case-control association test comparing the frequency of rare deleterious variants in PAH cases with population controls. The controls consisted of gnomAD WGS subjects as well unaffected parents from the Pediatric Cardiac Genomics Consortium (PCGC) (“internal controls”) ^45^ to increase statistical power. To control for confounding from genetic ancestry, we selected 1832 cases and 5262 internal controls of European ancestry using principle components analysis implemented in PLINK version 1.9 ^52^, and 7509 non-Finnish, European (NFE) gnomAD subjects. Relatedness was checked using Peddy ^52^, and only unrelated cases were included in the association test. To guard against batch effects in combined datasets from different sources ^53^, we applied heuristic filters including restriction of investigated regions to the intersection of regions captured by xGen, NimbleGen and MedExome; regions uniquely mapping to the reference sequence; regions with at least 10X coverage in 90% of samples; and excluding regions of low complexity. Variant filters included genotype quality >60, and removal of variants with allele fraction <0.25, alternate allele depth <4 or total read depth <10. We then tested for similarity of the rare synonymous variant rate among cases and controls.

The analysis of disease-associated genes was confined to gene-specific enrichment of rare, deleterious variants (AF ≤0.01%, LGD or D-mis). We used REVEL scores to predict the deleteriousness of missense variants. Instead of using a REVEL score threshold of 0.5 based on genome-wide variants^51^, we adapted a gene-specific variable threshold test ^54^. We performed binomial tests with variable REVEL score cut-offs ranging from 0.2-1 with 0.05 intervals, for each gene, and determined a cut-off is optimal if it achieves the smallest p-value (***P***_**0**_). Then we perform 10,000,000 permutations (shuffling of case-control label of rare genotypes); in each permutation, we obtained the smallest p-value (***P*’**) by the same variable REVEL threshold procedure. We define the empirical p-value of the gene as the fraction of permutations where ***P*’** is equal or smaller than ***P***_**0**_. We set REVEL score of LGD variants to 1 (more severe). In each binomial test, we assumed that under the null model, the number of rare deleterious variants observed in cases should follow a binomial distribution, given the total number of such variants in cases and controls, and a rate determined by fraction of cases in total number of subjects (cases and controls). We use binom.test function in R to calculate p-values in binomial tests. We defined the threshold for genome-wide significance by Bonferroni correction for multiple testing (n=20,000 genes, threshold p-value=2.5e−6). We used the Benjamini-Hochberg procedure to estimate false discovery rate (FDR) by p.adjust in R. All *GGCX* and *KLK1* variants reported herein were confirmed by Sanger sequencing.

## Acknowledgement

Samples and/or data from the National Biological Sample and Data Repository for PAH, which receives government support under an investigator-initiated grant (R24 HL105333 to WCN) awarded by the National Heart Lung and Blood Institute (NHLBI), were used in this study. We thank contributors, including the Pulmonary Hypertension Centers who collected samples used in this study, as well as patients and their families, whose help and participation made this work possible. Exome sequencing and genotyping data were generated by the Regeneron Genetics Center. Other funding support was provided by NHLBI HL060056 (WKC) and NIH R01GM120609 (YS).

## Authors’ contributions

MWP, WCN, WKC conceived and designed the study. WCN, MWP, WKC, LJM, HH, NZ, YS, JW analyzed and interpreted the data. WCN, MWP, CLW, NA, WKC, YS wrote the manuscript. KL, MWP, CW, JG, CLW, AWC, PAH Biobank collected patient samples and/or clinical information. CG-J provided and analyzed WES. All authors contributed and discussed the results and critically reviewed the manuscript.

