## supplemental data for "Novel risk genes and mechanisms implicated by exome sequencing of 2,572 individuals with pulmonary arterial hypertension"

**Supplementary Table 1. PAH Biobank cohort demographic data by PAH subclass.**

| | ALL | IPAH | APAH | | | | DTOX | FPAH | Other |
|---|---|---|---|---|---|---|---|---|---|
| | | | CTD | CHD | Portopulmonary | Other | | | |
| **Total, n (%)** | 2572 | 1110 (43.2) | 722 (28.1) | 268 (10.4) | 139 (5.4) | 110 (4.3) | 110 (4.3) | 98 (3.8) | 12 (0.5) |
| **Age-of-onset** | | | | | | | | | |
| Child (dx age <19) | 226 (8.8) | 94 (8.5) | 7 (0.9) | 95 (37.2)*** | 4 (2.8) | 6 (5.5) | 1 (0.9) | 13 (13.4) | 4 (33.3) |
| Adult (dx age >=19) | 2345 (91.2) | 1015 (91.4) | 715 (99.0) | 173 (62.8) | 135 (97.2) | 104 (94.5) | 109 (99.1) | 85 (86.6) | 8 (66.7) |
| **Gender** | | | | | | | | | |
| Female | 2023 (78.7) | 878 (78.4) | 655 (90.7) | 200 (74.5) | 63 (46.1) | 78 (70.9) | 81 (73.9) | 67 (68.2) | 9 (75.0) |
| Male | 548 (21.3) | 242 (21.6) | 67 (9.3) | 68 (25.5) | 76 (53.9) | 32 (29.1) | 29 (26.1) | 31 (31.6) | 3 (25.0) |
| Female:male ratio | 3.7:1 | 3.6:1 | 9.8:1 | 2.9:1 | 1:1.2 | 2.4:1 | 2.8:1 | 2.1:1 | 3:1 |
| **Ancestry n (%)** | | | | | | | | | |
| European | 1851 (72) | 816 (73) | 491 (68) | 190 (71) | 110 (79) | 63 (57.3) | 88 (80) | 87 (89) | 11 (91.7) |
| Hispanic | 316 (12) | 138 (12) | 74 (10) | 40 (15) | 25 (18)* | 18 (16.4) | 12 (11) | 9 (9.2) | 0 |
| African | 292 (11) | 118 (11) | 132 (18)* | 8 (3)** | 3 (2.2)** | 25 (22.7) | 5 (4.5) | 1 (1) | 1 (8.3) |
| East Asian | 70 (2.7) | 25 (2.2) | 20 (2.8) | 18 (6.7) | 1 (0.72) | 2 (1.8) | 4 (3.6) | 0 | 0 |
| South Asian | 28 (1.1) | 13 (1.2) | 4 (0.55) | 9 (3.4) | 0 | 2 (1.8) | 0 | 1 (1) | 0 |
| Others | 15 (0.58) | 10 (0.89) | 1 (0.14) | 3 (1.1) | 0 | 0 | 1 (0.91) | 0 | 0 |

APAH other included HIV, HHT and other rare associated diseases.
ALL other included 11 non-familial PVOD/PCH and one persistent pulmonary hypertension of the newborn.
*p=0.02, two-tailed Chi-square test (all Hispanic or African APAH cases vs Hispanic or African portopulmonary or CTD cases).
**p≤0.001, two-tailed Chi-square test (all African APAH cases vs African CHD or portopulmonary cases).
***<0.0001, two-tailed Chi-square test (all child-onset cases vs child-onset CHD cases).

**Supplementary Table 2. Rare, predicted deleterious variants* in established PAH risk genes among 2,572 PAH cases. Patients are heterozygous for the indicated variant unless noted.**

| ID | PAH subclass | Gender | Age dx | Ancestry | Gene | Transcript | Nucleotide change | Amino acid change | Variant type | Previously reported? | MAF (ExAC) | CADD | REVEL |
|---|---|---|---|---|---|---|---|---|---|---|---|---|---|
| 12-070 | APAH-HHT | F | 35 | EUR | *ACVRL1* | . | exon 10 del | . | exon deletion | no | N/A | . | . |
| 12-193 | APAH-HHT | F | 59 | EUR | *ACVRL1* | NM_001077401.1 | c.199C>T | p.(Arg67Trp) | D-Mis | [1-3] | . | 24 | 0.72 |
| 26-020 | DTOX | F | 35 | EUR | *ACVRL1* | NM_001077401.1 | c.430C>T | p.(Arg144*) | stopgain | [2] | 8.57E-06 | 39 | . |
| 07-010 | APAH-NF | F | 46 | EUR | *ACVRL1* | NM_001077401.1 | c.599G>A | p.(Arg200Gln) | D-Mis | rs1018788708 | . | 35 | 0.78 |
| 03-069 | APAH-CHD | F | 68 | EUR | *ACVRL1* | NM_001077401.1 | c.641G>T | p.(Gly214Val) | D-Mis | no | . | 28 | 0.83 |
| 08-069 | APAH-CTD | F | 53 | EUR | *ACVRL1* | NM_001077401.1 | c.721C>T | p.(Arg241Trp) | D-Mis | no | 5.09E-05 | 35 | 0.84 |
| 02-100* | FPAH | F | 55 | EUR | *ACVRL1* | NM_001077401.1 | c.841G>C | p.(Glu281Gln) | D-Mis | no | . | 30 | 0.81 |
| 12-015* | IPAH | F | 40 | EUR | *ACVRL1* | NM_001077401.1 | c.864T>G | p.(Phe288Leu) | D-Mis | no | . | 25 | 0.66 |
| 10-045 | APAH-HHT | F | 29 | EUR | *ACVRL1* | NM_001077401.1 | c.950T>C | p.(Ile317Thr) | D-Mis | [2, 4, 5] | . | 27 | 0.92 |
| 07-022 | APAH-CHD | M | 52 | EUR | *ACVRL1* | NM_001077401.1 | c.1064A>C | p.(His355Pro) | D-Mis | no | 8.39E-06 | 27 | 0.68 |
| 12-058 | APAH-HHT | F | 48 | EUR | *ACVRL1* | NM_001077401.1 | c.1120C>T | p.(Arg374Trp) | D-Mis | [1, 6] | . | 29 | 0.70 |
| 05-004 | IPAH | F | 46 | EUR | *ACVRL1* | NM_001077401.1 | c.1258G>A | p.(Asp420Asn) | D-Mis | no | . | 34 | 0.74 |
| 15-080 | IPAH | M | 14 | AMR | *ACVRL1* | NM_001077401.1 | c.1270C>A | p.(Pro424Thr) | D-Mis | [2, 4, 7] | . | 32 | 0.79 |
| 11-048 | APAH-HHT | F | 46 | EUR | *ACVRL1* | NM_001077401.1 | c.1331_1332dup | p.(Asp445Trpfs*21) | frameshift | no | . | 35 | . |
| 12-108 | APAH-HHT | F | 56 | SAS | *ACVRL1* | NM_001077401.1 | c.1345C>T | p.(Pro449Ser) | D-Mis | [1] | . | 33 | 0.74 |
| 01-018 | APAH-HHT | F | 4 | EUR | *ACVRL1* | NM_001077401.1 | c.1450C>T | p.(Arg484Trp) | D-Mis | [2, 6, 8, 9] | . | 33 | 0.92 |
| 22-065 | IPAH | F | 50 | EAS | *BMPR1A* | NM_004329.2 | c.407C>T | p.(Pro136Leu) | D-Mis | no | . | 27 | 0.83 |
| 22-110 | IPAH | F | 52 | AFR | *BMPR1A* | NM_004329.2 | c.1216C>T | p.(Arg406Cys) | D-Mis | rs587781332 | 2.47E-05 | 35 | 0.65 |
| 20-017 | APAH-HIV | F | 40 | AFR | *BMPR1A* | NM_004329.2 | c.1324C>T | p.(Arg442Cys) | D-Mis | rs587782496 | . | 35 | 0.74 |
| 06-130 | APAH-CHD | F | 27 | EUR | *BMPR1A* | NM_004329.2 | c.1498A>G | p.(Met500Val) | D-Mis | no | 4.94E-05 | 25 | 0.63 |
| 05-035 | APAH-CTD | M | 57 | AMR | *BMPR1B* | NM_001256792.1 | c.334C>T | p.(Pro112Ser) | D-Mis | no | . | 26 | 0.56 |
| 30-038 | APAH-Portopulm | F | 51 | AMR | *BMPR1B* | NM_001256792.1 | c.1253G>A | p.(Gly418Asp) | D-Mis | no | . | 33 | 0.77 |
| 04-012 | IPAH | M | 43 | EUR | *BMPR2* | NM_001204.6 | c.7del | p.(Ser3Profs*44) | frameshift | no | . | 24 | . |
| 04-026 | IPAH | F | 31 | EUR | *BMPR2* | NM_001204.6 | c.16C>T | p.(Gln6*) | stopgain | [2, 10-13] | . | 35 | . |
| 06-070 | IPAH | F | 28 | EUR | *BMPR2* | NM_001204.6 | c.16C>T | p.(Gln6*) | stopgain | [2, 10-13] | . | 35 | . |
| 08-097 | IPAH | M | 42 | EUR | *BMPR2* | NM_001204.6 | c.39G>A | p.(Trp13*) | stopgain | [2] | . | 36 | . |
| 02-072 | IPAH | F | 40 | AFR | *BMPR2* | NM_001204.6 | c.47G>A | p.(Trp16*) | stopgain | [2, 14, 15] | . | 35 | . |
| 06-027 | IPAH | F | 59 | EUR | *BMPR2* | NM_001204.6 | c.53_64del | p.(Ile18_Val21del) | in-frame | no | . | 18 | . |

| ID | PAH subclass | Gender | Age dx | Ancestry | Gene | Transcript | Nucleotide change | Amino acid change | Variant type | Previously reported? | MAF (ExAC) | CADD | REVEL |
|---|---|---|---|---|---|---|---|---|---|---|---|---|---|
| 15-067 | FPAH | F | 8 | SAS | *BMPR2* | NM_001204.6 | c.76+1G>T | p.(=) | splicing | [2] | . | 23 | . |
| 08-054 | FPAH | F | 37 | EUR | *BMPR2* | NM_001204.6 | c.76+2T>C | p.(=) | splicing | [2] | . | 23 | . |
| 11-049 | IPAH | F | 33 | EUR | *BMPR2* | NM_001204.6 | c.116del | p.(Pro39Argfs*8) | frameshift | no | . | 33 | . |
| 10-096 | IPAH | F | 60 | EUR | *BMPR2* | NM_001204.6 | c.118dup | p.(Tyr40Leufs*9) | frameshift | no | . | 32 | . |
| 02-071* | FPAH | F | 40 | EUR | *BMPR2* | NM_001204.6 | c.135dup | p.(Ile46Aspfs*3) | frameshift | no | . | . | . |
| 17-077 | IPAH | F | 38 | AMR | *BMPR2* | NM_001204.6 | c.164dup | p.(Asn55Lysfs*10) | frameshift | no | . | 33 | . |
| 22-034 | IPAH | F | 36 | AFR | *BMPR2* | NM_001204.6 | c.199dup | p.(Tyr67Leufs*31) | frameshift | no | . | 33 | . |
| 12-115 | FPAH | M | 28 | EUR | *BMPR2* | NM_001204.6 | c.200A>G | p.(Tyr67Cys) | D-Mis | [2, 6, 8, 13, 16-18] | . | 26 | 0.89 |
| 06-010 | FPAH | M | 56 | EUR | *BMPR2* | NM_001204.6 | c.201T>G | p.(Tyr67*) | stopgain | no | . | 35 | . |
| 06-011 | FPAH | F | 41 | EUR | *BMPR2* | NM_001204.6 | c.201T>G | p.(Tyr67*) | stopgain | no | . | 35 | . |
| 17-002 | IPAH | F | 34 | EAS | *BMPR2* | NM_001204.6 | c.203del | (p.(Gly68Alafs*10) | frameshift | no | . | 33 | . |
| 09-059 | IPAH | F | 44 | EUR | *BMPR2* | NM_001204.6 | c.211del | p.(Glu71Argfs*7) | frameshift | no | . | 33 | . |
| 06-116 | APAH-CHD | F | 4 | EUR | *BMPR2* | NM_001204.6 | c.211G>A | p.(Glu71Lys) | D-Mis | [19] | . | 26 | 0.75 |
| 27-004 | IPAH | F | 33 | EUR | *BMPR2* | NM_001204.6 | c.218C>A | p.(Ser73*) | stopgain | no | . | 36 | . |
| 05-094 | IPAH | F | 24 | AMR | *BMPR2* | NM_001204.6 | c.251G>A | p.(Cys84Tyr) | D-Mis | [8] | . | 28 | 0.95 |
| 22-054 | IPAH | F | 37 | EUR | *BMPR2* | NM_001204.6 | c.255G>A | p.(Trp85*) | stopgain | [2, 10, 18] | . | 38 | . |
| 32-007 | APAH-CTD | F | 68 | EUR | *BMPR2* | NM_001204.6 | c.255G>A | p.(Trp85*) | stopgain | [2, 10, 18] | . | 38 | . |
| 03-003 | IPAH | M | 49 | EUR | *BMPR2* | NM_001204.6 | c.258del | p.(His87Thrfs*14) | frameshift | no | . | 33 | . |
| 08-048 | IPAH | M | 44 | EUR | *BMPR2* | NM_001204.6 | c.277dup | p.(Glu93Glyfs*5) | frameshift | no | . | 34 | . |
| 04-007 | IPAH | F | 21 | EUR | *BMPR2* | NM_001204.6 | c.278_279del | p.(Glu93Valfs*4) | frameshift | no | . | 34 | . |
| 15-051* | IPAH | F | 4 | EUR | *BMPR2* | NM_001204.6 | c.295T>C | p.(Cys99Arg) | D-Mis | [2, 6, 18, 20, 21] | . | 26 | 0.94 |
| 06-022 | IPAH | M | 3 | EUR | *BMPR2* | NM_001204.6 | c.297T>G | p.(Cys99Trp) | D-Mis | [6] | . | 26 | 0.93 |
| 16-032 | IPAH | F | 21 | EUR | *BMPR2* | NM_001204.6 | c.344del | p.(Phe115Serfs*37) | frameshift | no | . | 33 | . |
| 16-050 | FPAH | F | 41 | EUR | *BMPR2* | NM_001204.6 | c.344del | p.(Phe115Serfs*37) | frameshift | no | . | 33 | . |
| 12-088 | FPAH | F | 29 | EUR | *BMPR2* | NM_001204.6 | c.350G>A | p.(Cys117Tyr) | D-Mis | [2, 6, 21, 22] | . | 28 | 0.96 |
| 19-032 | FPAH | F | 28 | EUR | *BMPR2* | NM_001204.6 | c.350G>A | p.(Cys117Tyr) | D-Mis | [2, 6, 21, 22] | . | 28 | 0.96 |
| 19-010 | IPAH | M | 33 | EUR | *BMPR2* | NM_001204.6 | c.353G>A | p.(Cys118Tyr) | D-Mis | [2, 23, 24] | . | 28 | 0.94 |
| 19-003 | FPAH | F | 28 | EUR | *BMPR2* | NM_001204.6 | c.354T>G | p.(Cys118Trp) | D-Mis | [2, 25] | . | 26 | 0.93 |
| 19-070 | IPAH | F | 30 | EUR | *BMPR2* | NM_001204.6 | c.354T>G | p.(Cys118Trp) | D-Mis | [2, 25] | . | 26 | 0.93 |
| 19-077 | FPAH | F | 39 | EUR | *BMPR2* | NM_001204.6 | c.354T>G | p.(Cys118Trp) | D-Mis | [2, 25] | . | 26 | 0.93 |

| ID | PAH subclass | Gender | Age dx | Ancestry | Gene | Transcript | Nucleotide change | Amino acid change | Variant type | Previously reported? | MAF (ExAC) | CADD | REVEL |
|---|---|---|---|---|---|---|---|---|---|---|---|---|---|
| 19-088 | FPAH | F | 26 | EUR | *BMPR2* | NM_001204.6 | c.354T>G | p.(Cys118Trp) | D-Mis | [2, 25] | . | 26 | 0.93 |
| 25-001 | FPAH | M | 3 | EUR | *BMPR2* | NM_001204.6 | c.354T>G | p.(Cys118Trp) | D-Mis | [2, 25] | . | 26 | 0.93 |
| 12-095 | IPAH | M | 45 | EUR | *BMPR2* | NM_001204.6 | c.367T>C | p.(Cys123Arg) | D-Mis | [2, 7, 8, 16, 23] | . | 27 | 0.93 |
| 12-101 | IPAH | M | 64 | EUR | *BMPR2* | NM_001204.6 | c.367T>C | p.(Cys123Arg) | D-Mis | [2, 7, 8, 16, 23] | . | 27 | 0.93 |
| 12-197 | FPAH | M | 39 | EUR | *BMPR2* | NM_001204.6 | c.377A>G | p.(Asn126Ser) | D-Mis | [2, 8, 12, 16, 21, 24, 26] | . | 26 | 0.80 |
| 19-062 | IPAH | F | 48 | EUR | *BMPR2* | NM_001204.6 | c.377A>G | p.(Asn126Ser) | D-Mis | [2, 8, 12, 16, 21, 24, 26] | . | 26 | 0.80 |
| 13-045 | DTOX | M | 26 | EUR | *BMPR2* | NM_001204.6 | c.419-1G>T | p.(=) | splicing | no | . | 26 | . |
| 02-100* | FPAH | F | 55 | EUR | *BMPR2* | NM_001204.6 | c.439C>T | p.(Arg147*) | stopgain | [2, 6, 8, 13, 16-18, 20, 23, 24, 26-28] | . | 38 | . |
| 20-011 | IPAH | F | 36 | EUR | *BMPR2* | NM_001204.6 | c.439C>T | p.(Arg147*) | stopgain | [2, 6, 8, 13, 16-18, 20, 23, 24, 26-28] | . | 38 | . |
| 26-029 | IPAH | F | 21 | EUR | *BMPR2* | NM_001204.6 | c.452_453del | p.(Ile151Asnfs*29) | stopgain | no | . | 34 | . |
| 26-039 | IPAH | F | 31 | EUR | *BMPR2* | NM_001204.6 | c.524del | p.(Leu175*) | stopgain | no | . | 34 | . |
| 03-097 | IPAH | F | 31 | EUR | *BMPR2* | NM_001204.6 | c.529+1G>A | p.(=) | splicing | [19] | . | 28 | . |
| 18-038 | IPAH | F | 54 | EUR | *BMPR2* | NM_001204.6 | c.529+1G>A | p.(=) | splicing | [19] | . | 28 | . |
| 02-119 | IPAH | F | 53 | EUR | *BMPR2* | NM_001204.6 | c.529G>A | p.(Gly177Arg) | D-Mis | no | . | 23 | 0.55 |
| 18-078 | IPAH | F | 21 | AFR | *BMPR2* | NM_001204.6 | c.543_544del | p.(Gly182Serfs*17) | frameshift | no | . | 35 | . |
| 03-110 | IPAH | F | 36 | AFR | *BMPR2* | NM_001204.6 | c.631C>T | p.(Arg211*) | stopgain | [2, 6, 8, 18 16, 20-24, 29, 30] | . | 42 | . |
| 21-054 | IPAH | F | 48 | EUR | *BMPR2* | NM_001204.6 | c.631C>T | p.(Arg211*) | stopgain | [2, 6, 8, 18 16, 20-24, 29, 30] | . | 42 | . |
| 22-001 | IPAH | F | 55 | EUR | *BMPR2* | NM_001204.6 | c.631C>T | p.(Arg211*) | stopgain | [2, 6, 8, 18 16, 20-24, 29, 30] | . | 42 | . |
| 06-015 | FPAH | F | 60 | EUR | *BMPR2* | NM_001204.6 | c.637C>T | p.(Arg213*) | stopgain | [2, 8, 13, 18, 20] | . | 37 | . |
| 10-012 | FPAH | F | 30 | EUR | *BMPR2* | NM_001204.6 | c.637C>T | p.(Arg213*) | stopgain | [2, 8, 13, 18, 20] | . | 37 | . |
| 12-015* | IPAH | F | 40 | EUR | *BMPR2* | NM_001204.6 | c.637C>T | p.(Arg213*) | stopgain | [2, 8, 13, 18, 20] | . | 37 | . |
| 26-037 | FPAH | F | 26 | EUR | *BMPR2* | NM_001204.6 | c.637C>T | p.(Arg213*) | stopgain | [2, 8, 13, 18, 20] | . | 37 | . |
| 16-061 | IPAH | F | 37 | Dominican | *BMPR2* | NM_001204.6 | c.688A>T | p.(Lys230*) | stopgain | no | . | 40 | . |
| 16-041 | IPAH | F | 42 | EUR | *BMPR2* | NM_001204.6 | c.689_690del | p.(Lys230Serfs*25) | frameshift | no | . | 34 | . |
| 08-074 | FPAH | F | 19 | EUR | *BMPR2* | NM_001204.6 | c.697del | p.(Cys397*) | stopgain | no | . | 35 | . |
| 10-100 | IPAH | M | 34 | EUR | *BMPR2* | NM_001204.6 | c.712C>T | p.(Gln238*) | stopgain | no | . | 39 | . |
| 07-011 | APAH-CTD | F | 65 | EUR | *BMPR2* | NM_001204.6 | c.775C>T | p.(Arg259Cys) | D-Mis | no | 4.95E-05 | 34 | 0.78 |
| 12-208 | APAH-unspecified | F | 38 | AMR | *BMPR2* | NM_001204.6 | c.797G>C | p.(Arg266Thr) | D-Mis | [2, 18, 21] | 4.13E-05 | 28 | 0.63 |
| 05-046 | IPAH | F | 32 | EUR | *BMPR2* | NM_001204.6 | c.834dup | p.(Met279Aspfs*19) | frameshift | no | . | 35 | . |
| 07-060 | FPAH | F | 34 | EUR | *BMPR2* | NM_001204.6 | c.846T>G | p.(Tyr282*) | stopgain | rs863223419 | . | 36 | . |

| ID | PAH subclass | Gender | Age dx | Ancestry | Gene | Transcript | Nucleotide change | Amino acid change | Variant type | Previously reported? | MAF (ExAC) | CADD | REVEL |
|---|---|---|---|---|---|---|---|---|---|---|---|---|---|
| 03-058 | IPAH | F | 39 | EUR | *BMPR2* | NM_001204.6 | c.852+1G>A | p.(=) | splicing | [2, 16] | . | 27 | . |
| 19-014 | FPAH | M | 37 | EUR | *BMPR2* | NM_001204.6 | c.852+1G>A | p.(=) | splicing | [2, 16] | . | 27 | . |
| 19-019 | FPAH | F | 48 | EUR | *BMPR2* | NM_001204.6 | c.852+1G>A | p.(=) | splicing | [2, 16] | . | 27 | . |
| 05-008 | IPAH | M | 40 | EUR | *BMPR2* | NM_001204.6 | c.853-1G>A | p.(=) | splicing | [2, 8] | . | 26 | . |
| 07-077 | FPAH | M | 43 | EUR | *BMPR2* | NM_001204.6 | c.853-2A>G | p.(=) | splicing | [2, 8, 10, 11, 18] | . | 25 | . |
| 08-104 | FPAH | F | 26 | EUR | *BMPR2* | NM_001204.6 | c.862dup | p.(Cys288Leufs*10) | frameshift | no | . | 33 | . |
| 05-052 | APAH-CHD | F | 32 | AMR | *BMPR2* | NM_001204.6 | c.872dup | p.(Leu291Phefs*7) | frameshift | no | . | 35 | . |
| 08-032 | IPAH | F | 51 | EUR | *BMPR2* | NM_001204.6 | c.893G>A | p.(Trp298*) | stopgain | [2, 31] | . | 37 | . |
| 19-017 | FPAH | M | 33 | EUR | *BMPR2* | NM_001204.6 | c.894_895dup | p.(Val299Glyfs*2) | frameshift | no | . | 35 | . |
| 28-136 | IPAH | F | 32 | EUR | *BMPR2* | NM_001204.6 | c.918_921del | p.(His306Glnfs*28) | frameshift | no | . | 35 | . |
| 07-096 | IPAH | F | 28 | AMR | *BMPR2* | NM_001204.6 | c.935T>C | p.(Leu312Pro) | D-Mis | no | . | 29 | 0.98 |
| 16-017 | IPAH | F | 41 | AMR | *BMPR2* | NM_001204.6 | c.942_943insA | p.(Leu315Thrfs*12) | frameshift | no | . | 35 | . |
| 17-049 | IPAH | F | 35 | EUR | *BMPR2* | NM_001204.6 | c.947A>G | p.(His316Arg) | D-Mis | no | . | 25 | 0.93 |
| 11-018 | APAH-CTD | F | 33 | EUR | *BMPR2* | NM_001204.6 | c.961C>T | p.(Arg321*) | stopgain | [2, 6, 8, 16-18, 24, 32] | . | 40 | . |
| 13-083 | FPAH | F | 62 | EUR | *BMPR2* | NM_001204.6 | c.961C>T | p.(Arg321*) | stopgain | [2, 6, 8, 16-18, 24, 32] | . | 40 | . |
| 17-027 | IPAH | F | 31 | EUR | *BMPR2* | NM_001204.6 | c.961C>T | p.(Arg321*) | stopgain | [2, 6, 8, 16-18, 24, 32] | . | 40 | . |
| 18-043 | IPAH | M | 30 | AMR | *BMPR2* | NM_001204.6 | c.961C>T | p.(Arg321*) | stopgain | [2, 6, 8, 16-18, 24, 32] | . | 40 | . |
| 37-011 | IPAH | M | 69 | EUR | *BMPR2* | NM_001204.6 | c.961C>T | p.(Arg321*) | stopgain | [2, 6, 8, 16-18, 24, 32] | . | 40 | . |
| 15-077 | IPAH | F | 16 | EUR | *BMPR2* | NM_001204.6 | c.967G>A | p.(Asp323Asn) | D-Mis | no | . | 32 | 0.60 |
| 12-084 | FPAH | M | 45 | EUR | *BMPR2* | NM_001204.6 | c.969dup | p.(His324Serfs*3) | frameshift | no | . | 34 | . |
| 05-027 | IPAH | M | 37 | EUR | *BMPR2* | NM_001204.6 | c.994C>T | p.(Arg332*) | stopgain | [6, 8, 16, 20, 22, 23, 26, 30, 33, 34] | . | 38 | . |
| 08-082 | IPAH | F | 35 | EUR | *BMPR2* | NM_001204.6 | c.994C>T | p.(Arg332*) | stopgain | [6, 8, 16, 20, 22, 23, 26, 30, 33, 34] | . | 38 | . |
| 10-018 | FPAH | M | 54 | EUR | *BMPR2* | NM_001204.6 | c.994C>T | p.(Arg332*) | stopgain | [6, 8, 16, 20, 22, 23, 26, 30, 33, 34] | . | 38 | . |
| 19-002 | FPAH | M | 26 | EUR | *BMPR2* | NM_001204.6 | c.994C>T | p.(Arg332*) | stopgain | [6, 8, 16, 20, 22, 23, 26, 30, 33, 34] | . | 38 | . |
| 19-021 | FPAH | F | 14 | EUR | *BMPR2* | NM_001204.6 | c.994C>T | p.(Arg332*) | stopgain | [6, 8, 16, 20, 22, 23, 26, 30, 33, 34] | . | 38 | . |
| 25-009 | FPAH | M | 9 | EUR | *BMPR2* | NM_001204.6 | c.994C>T | p.(Arg332*) | stopgain | [6, 8, 16, 20, 22, 23, 26, 30, 33, 34] | . | 38 | . |
| 02-045 | FPAH | F | 48 | EUR | *BMPR2* | NM_001204.6 | c.995G>C | p.(Arg332Pro) | D-Mis | [6] | . | 34 | 0.99 |
| 06-013 | IPAH | F | 31 | EUR | *BMPR2* | NM_001204.6 | c.1040G>A | p.(Cys347Tyr) | D-Mis | [2, 6, 18, 25] | . | 29 | 0.94 |
| 19-090 | IPAH | F | 53 | EUR | *BMPR2* | NM_001204.6 | c.1126G>T | p.(Glu376*) | stopgain | [2] | . | 45 | . |
| 10-031 | FPAH | F | 22 | EUR | *BMPR2* | NM_001204.6 | c.1128del | p.(Val377Leufs*12) | frameshift | no | . | 26 | . |

| ID | PAH subclass | Gender | Age dx | Ancestry | Gene | Transcript | Nucleotide change | Amino acid change | Variant type | Previously reported? | MAF (ExAC) | CADD | REVEL |
|---|---|---|---|---|---|---|---|---|---|---|---|---|---|
| 10-032 | FPAH | F | 53 | EUR | *BMPR2* | NM_001204.6 | c.1128del | p.(Val377Leufs*12) | frameshift | no | . | 26 | . |
| 05-136 | FPAH | F | 22 | AMR | *BMPR2* | NM_001204.6 | c.1128+1G>A | p.(=) | splicing | [2, 18] | . | 27 | . |
| 06-007 | IPAH | M | 26 | EUR | *BMPR2* | NM_001204.6 | c.1128+1G>A | p.(=) | splicing | [2, 18] | . | 27 | . |
| 15-059 | FPAH | F | 16 | AMR | *BMPR2* | NM_001204.6 | c.1128+1G>A | p.(=) | splicing | [2, 18] | . | 27 | . |
| 03-113 | IPAH | M | 30 | EUR | *BMPR2* | NM_001204.6 | c.1128+1G>C | p.(=) | splicing | [2, 18] | . | 26 | . |
| 07-075 | FPAH | F | 35 | EUR | *BMPR2* | NM_001204.6 | c.1141dup | p.(Arg381Lysfs*18) | frameshift | no | . | 34 | . |
| 06-059 | IPAH | F | 7 | EAS | *BMPR2* | NM_001204.6 | c.1154C>G | p.(Pro385Arg) | D-Mis | [6] | . | 28 | 0.98 |
| 04-008 | IPAH | M | 34 | AFR | *BMPR2* | NM_001204.6 | c.1172C>A | p.(Ala391Asp) | D-Mis | no | . | 33 | 0.82 |
| 25-005 | APAH-CHD | F | 10 | EUR | *BMPR2* | NM_001204.6 | c.1175T>A | p.(Val392Glu) | D-Mis | no | . | 32 | 0.91 |
| 09-038 | IPAH | F | 27 | AFR | *BMPR2* | NM_001204.6 | c.1191_1192del | p.(Cys397*) | stopgain | no | . | 35 | . |
| 10-049 | IPAH | F | 57 | EUR | *BMPR2* | NM_001204.6 | c.1197del | p.(Ala400Leufs*2) | frameshift | no | . | 35 | . |
| 13-035 | FPAH | F | 35 | AFR | *BMPR2* | NM_001204.6 | c.1233_1236dup | p.(Tyr413Asnfs*36) | frameshift | no | . | 35 | . |
| 17-013 | FPAH | F | 38 | EUR | *BMPR2* | NM_001204.6 | c.1250_1253del | p.(Phe417*) | stopgain | no | . | 34 | . |
| 08-037 | IPAH | F | 67 | EUR | *BMPR2* | NM_001204.6 | c.1361C>T | p.(Ser454Phe) | D-Mis | no | . | 33 | 0.52 |
| 12-085 | IPAH | F | 63 | EUR | *BMPR2* | NM_001204.6 | c.1361C>T | p.(Ser454Phe) | D-Mis | no | . | 33 | 0.52 |
| 10-005 | FPAH | F | 44 | EUR | *BMPR2* | NM_001204.6 | c.1397G>A | p.(Trp466*) | stopgain | [18, 32 20, 24] | . | 43 | . |
| 08-022 | IPAH | F | 60 | EUR | *BMPR2* | NM_001204.6 | c.1413+1G>A | p.(=) | splicing | [2, 16, 24] | . | 25 | . |
| 19-009 | IPAH | F | 29 | EUR | *BMPR2* | NM_001204.6 | c.1450T>C | p.(Trp484Arg) | D-Mis | no | . | 28 | 0.79 |
| 08-003 | FPAH | F | 48 | EUR | *BMPR2* | NM_001204.6 | c.1451G>A | p.(Trp484*) | stopgain | [2] | . | 40 | . |
| 08-068 | FPAH | F | 30 | EUR | *BMPR2* | NM_001204.6 | c.1451G>A | p.(Trp484*) | stopgain | [2] | . | 40 | . |
| 33-009 | IPAH | F | 25 | AFR | *BMPR2* | NM_001204.6 | c.1451G>A | p.(Trp484*) | stopgain | [2] | . | 40 | . |
| 33-012 | IPAH | F | 58 | AFR | *BMPR2* | NM_001204.6 | c.1451G>A | p.(Trp484*) | stopgain | [2] | . | 40 | . |
| 04-083 | IPAH | F | 40 | EUR | *BMPR2* | NM_001204.6 | c.1460A>T | p.(Asp487Val) | D-Mis | [2, 21, 24] | . | 29 | 0.95 |
| 05-194 | IPAH | F | 61 | EUR | *BMPR2* | NM_001204.6 | c.1468G>A | p.(Ala490Thr) | D-Mis | no | . | 33 | 0.52 |
| 39-001 | IPAH | F | 41 | AFR | *BMPR2* | NM_001204.6 | c.1471C>T | p.(Arg491Trp) | D-Mis | [2, 16-18, 32, 33, 35-37 6, 8, 11, 12, 24] | . | 35 | 0.96 |
| 03-108 | FPAH | F | 30 | EUR | *BMPR2* | NM_001204.6 | c.1472G>A | p.(Arg491Gln) | D-Mis | [2, 5, 6, 8, 18 24, 26, 34, 35] | . | 35 | 0.96 |
| 06-047 | IPAH | M | 11 | AMR | *BMPR2* | NM_001204.6 | c.1472G>A | p.(Arg491Gln) | D-Mis | [2, 5, 6, 8, 18 24, 26, 34, 35] | . | 35 | 0.96 |
| 08-020 | IPAH | M | 52 | EUR | *BMPR2* | NM_001204.6 | c.1472G>A | p.(Arg491Gln) | D-Mis | [2, 5, 6, 8, 18 24, 26, 34, 35] | . | 35 | 0.96 |
| 19-018 | IPAH | M | 43 | EUR | *BMPR2* | NM_001204.6 | c.1472G>A | p.(Arg491Gln) | D-Mis | [2, 5, 6, 8, 18 24, 26, 34, 35] | . | 35 | 0.96 |
| 02-155 | IPAH | F | 13 | AMR | *BMPR2* | NM_001204.6 | c.1483C>T | p.(Gln495*) | stopgain | [2, 24, 32] | . | 41 | . |

| ID | PAH subclass | Gender | Age dx | Ancestry | Gene | Transcript | Nucleotide change | Amino acid change | Variant type | Previously reported? | MAF (ExAC) | CADD | REVEL |
|---|---|---|---|---|---|---|---|---|---|---|---|---|---|
| 03-107 | APAH-CTD | F | 76 | EUR | *BMPR2* | NM_001204.6 | c.1492G>A | p.(Glu498Lys) | D-Mis | no | . | 34 | 0.70 |
| 29-006 | FPAH | F | 36 | EUR | *BMPR2* | NM_001204.6 | c.1524G>A | p.(Trp508*) | stopgain | [2, 24] | . | 40 | . |
| 08-060 | FPAH | M | 36 | EUR | *BMPR2* | NM_001204.6 | c.1549del | p.(Thr517Glnfs*47) | frameshift | No | . | 35 | . |
| 22-060 | IPAH | F | 41 | EUR | *BMPR2* | NM_001204.6 | c.1744A>T | p.(Lys582*) | stopgain | No | . | 42 | . |
| 02-058 | IPAH | F | 68 | AMR | *BMPR2* | NM_001204.6 | c.1748dup | p.(Asn583Lysfs*6) | frameshift | No | . | 35 | . |
| 26-015 | APAH-unspecified | F | 51 | EUR | *BMPR2* | NM_001204.6 | c.1939_1940del | p.(Gln647Valfs*27) | frameshift | No | . | 34 | . |
| 26-023 | IPAH | F | 44 | EUR | *BMPR2* | NM_001204.6 | c.1958del | p.(Pro653Leufs*6) | frameshift | [2, 13, 38] | . | 29 | . |
| 13-079 | IPAH | F | 54 | EUR | *BMPR2* | NM_001204.6 | c.1981G>T | p.(Glu661*) | stopgain | [2] | . | 43 | . |
| 22-015 | IPAH | F | 49 | AFR | *BMPR2* | NM_001204.6 | c.2073dup | p.(Gln692Thrfs*12) | frameshift | no | . | 33 | . |
| 13-050 | APAH-HIV | F | 39 | AFR | *BMPR2* | NM_001204.6 | c.2140G>T | p.(Ala714Ser) | D-Mis | no | 9.08E-05 | 24 | 0.59 |
| 08-027 | IPAH | F | 30 | AFR | *BMPR2* | NM_001204.6 | c.2146G>T | p.(Glu716*) | stopgain | no | . | 45 | . |
| 04-001 | IPAH | F | 36 | EUR | *BMPR2* | NM_001204.6 | c.2202del | p.(Pro735Leufs*26) | frameshift | no | . | 31 | . |
| 12-152 | FPAH | F | 40 | EUR | *BMPR2* | NM_001204.6 | c.2216del | p.(Pro739Leufs*22) | frameshift | no | . | 35 | . |
| 07-086 | IPAH | F | 47 | EUR | *BMPR2* | NM_001204.6 | c.2268del | p.(Ser757Valfs*4) | frameshift | no | . | 35 | . |
| 06-008 | APAH-CHD | M | 2 | EUR | *BMPR2* | NM_001204.6 | c.2353G>A | p.(Glu785Lys) | D-Mis | [19] | 8.24E-06 | 32 | 0.59 |
| 26-033 | IPAH | M | 66 | EUR | *BMPR2* | NM_001204.6 | c.2396dup | p.(His800Serfs*13) | frameshift | no | . | 34 | . |
| 03-042 | FPAH | M | 54 | EUR | *BMPR2* | NM_001204.6 | c.2410_2413del | p.(Val804Profs*2) | frameshift | no | . | 35 | . |
| 08-062 | FPAH | F | 30 | EUR | *BMPR2* | NM_001204.6 | c.2450_2451del | p.(Asn817Ilefs*25) | frameshift | no | . | 35 | . |
| 08-034 | FPAH | M | 13 | EUR | *BMPR2* | NM_001204.6 | c.2450_2451del | p.(Asn817Ilefs*25) | frameshift | no | . | 35 | . |
| 08-035 | FPAH | F | 46 | EUR | *BMPR2* | NM_001204.6 | c.2450_2451del | p.(Asn817Ilefs*25) | frameshift | no | . | 35 | . |
| 02-011 | DTOX | F | 55 | EUR | *BMPR2* | NM_001204.6 | c.2457_2464del | p.(Ala820Asnfs*20) | frameshift | no | . | 35 | . |
| 12-048 | FPAH | F | 29 | EUR | *BMPR2* | NM_001204.6 | c.2580del | p.(Asn861Ilefs*11) | frameshift | no | . | 34 | . |
| 10-083 | FPAH | F | 20 | EUR | *BMPR2* | NM_001204.6 | c.2617C>T | p.(Arg873*) | stopgain | [2, 8, 18 20, 24, 26, 28, 35, 36] | . | 44 | . |
| 13-059 | IPAH | F | 26 | EUR | *BMPR2* | NM_001204.6 | c.2617C>T | p.(Arg873*) | stopgain | [2, 8, 18 20, 24, 26, 28, 35, 36] | . | 44 | . |
| 02-099 | DTOX | F | 38 | AMR | *BMPR2* | NM_001204.6 | c.2651_2652insAT | p.(Asp885Trpfs*12) | frameshift | no | . | 34 | . |
| 02-148 | IPAH | F | 28 | AFR | *BMPR2* | NM_001204.6 | c.2695C>T | p.(Arg899*) | stopgain | [2, 8, 14, 16, 17, 21, 24, 26] | . | 39 | . |
| 08-019 | IPAH | F | 35 | EUR | *BMPR2* | NM_001204.6 | c.2695C>T | p.(Arg899*) | stopgain | [2, 8, 21, 24 16 14, 17, 26] | . | 39 | . |
| 14-033 | FPAH | F | 40 | EUR | *BMPR2* | NM_001204.6 | c.2695C>T | p.(Arg899*) | stopgain | [2, 8, 14, 16, 17, 21, 24, 26] | . | 39 | . |
| 28-008 | IPAH | F | 41 | EUR | *BMPR2* | NM_001204.6 | c.2695C>T | p.(Arg899*) | stopgain | [2, 8, 14, 16, 17, 21, 24, 26] | . | 39 | . |
| 03-051 | IPAH | F | 41 | EUR | *BMPR2* | NM_001204.6 | c.2730T>A | p.(Cys910*) | stopgain | [2] | . | 36 | . |

| ID | PAH subclass | Gender | Age dx | Ancestry | Gene | Transcript | Nucleotide change | Amino acid change | Variant type | Previously reported? | MAF (ExAC) | CADD | REVEL |
|---|---|---|---|---|---|---|---|---|---|---|---|---|---|
| 14-044 | IPAH | F | 53 | AMR | *BMPR2* | NM_001204.6 | c.2952del | p.(Trp984Cysfs*50) | frameshift | no | . | 35 | . |
| 10-021 | IPAH | F | 55 | EUR | *BMPR2* | . | exon1 del | . | exon deletion | [23, 39] | N/A | . | . |
| 11-010 | IPAH | F | 26 | EUR | *BMPR2* | . | exon 1 del | . | exon deletion | [23, 39] | N/A | . | . |
| 03-029 | FPAH | M | 33 | EUR | *BMPR2* | . | exon 2-3 del | . | exon deletion | [40] | N/A | . | . |
| 04-019 | IPAH | F | 44 | EUR | *BMPR2* | . | exon 2-3 del | . | exon deletion | [40] | N/A | . | . |
| 04-075 | IPAH | F | 22 | EUR | *BMPR2* | . | exon 2-3 del | . | exon deletion | [40] | N/A | . | . |
| 10-010 | IPAH | F | 21 | EUR | *BMPR2* | . | exon 2-3 del | . | exon deletion | [40] | N/A | . | . |
| 12-207 | IPAH | F | 35 | EUR | *BMPR2* | . | exon 2-3 del | . | exon deletion | [40] | N/A | . | . |
| 04-060 | IPAH | M | 55 | EUR | *BMPR2* | . | exon 3 del | . | exon deletion | [41] | N/A | . | . |
| 08-023 | FPAH | F | 47 | EUR | *BMPR2* | . | exon 3 del | . | exon deletion | [41] | N/A | . | . |
| 08-073 | FPAH | M | 58 | EUR | *BMPR2* | . | exon 3 del | . | exon deletion | [41] | N/A | . | . |
| 08-114 | FPAH | F | 34 | EUR | *BMPR2* | . | exon 3 del | . | exon deletion | [41] | N/A | . | . |
| 12-013 | FPAH | F | 59 | EUR | *BMPR2* | . | exon 3 del | . | exon deletion | [41] | N/A | . | . |
| 10-095 | FPAH | M | 69 | EUR | *BMPR2* | . | exon 4 del | . | exon deletion | no | N/A | . | . |
| 22-017 | FPAH | F | 45 | EUR | *BMPR2* | . | exon 4 del | . | exon deletion | no | N/A | . | . |
| 07-006 | IPAH | F | 42 | EUR | *BMPR2* | . | exon 4-5 del | . | exon deletion | [41, 42] | N/A | . | . |
| 05-152 | IPAH | F | 36 | AMR | *BMPR2* | . | exon 4-7 del | . | exon deletion | no | N/A | . | . |
| 10-002 | FPAH | M | 26 | EUR | *BMPR2* | . | exon 4-7 del | . | exon deletion | no | N/A | . | . |
| 14-039 | FPAH | M | 60 | EUR | *BMPR2* | . | exon 4-7 del | . | exon deletion | no | N/A | . | . |
| 05-109 | IPAH | F | 22 | AMR | *BMPR2* | . | exon 4-8 del | . | exon deletion | no | N/A | . | . |
| 26-038 | IPAH | F | 64 | EUR | *BMPR2* | . | exon 4-9 del | . | exon deletion | no | N/A | . | . |
| 08-026 | IPAH | F | 56 | EUR | *BMPR2* | . | exon 6 del | . | exon deletion | [16] | N/A | . | . |
| 02-139 | FPAH | F | 25 | AMR | *BMPR2* | . | exon 8-9 del | . | exon deletion | [39] | N/A | . | . |
| 08-096 | IPAH | F | 32 | EUR | *BMPR2* | . | exon 11-12 del | . | exon deletion | [23, 39] | N/A | . | . |
| 17-015 | APAH-Portopulm | M | 56 | EAS | *CAV1* | NM_001172896.1 | c.-13_-10del | p.(=) | splicing | no | 1.65E-05 | 24 | . |
| 17-012 | IPAH | F | 33 | AFR | *CAV1* | NM_001172896.1 | c.191C>T | p.(Thr64Met) | D-Mis | no | 2.49E-05 | 31 | 0.92 |
| 04-038 | IPAH | F | 29 | EUR | *CAV1* | NM_001172896.1 | c.209G>A | p.(Arg70His) | D-Mis | no | 3.31E-05 | 32 | 0.96 |
| 17-064 | APAH-CHD | F | 54 | EUR | *CAV1* | NM_001172896.1 | c.274del | p.(Ser92Leufs*16) | frameshift | no | . | 35 | . |
| 08-041 | FPAH | F | 5 | EUR | *CAV1* | NM_001172896.1 | c.381del | p.(Leu128Serfs*22) | frameshift | no | . | 32 | . |

| ID | PAH subclass | Gender | Age dx | Ancestry | Gene | Transcript | Nucleotide change | Amino acid change | Variant type | Previously reported? | MAF (ExAC) | CADD | REVEL |
|---|---|---|---|---|---|---|---|---|---|---|---|---|---|
| 08-086 | FPAH | F | 41 | EUR | *CAV1* | NM_001172896.1 | c.381del | p.(Leu128Serfs*22) | frameshift | no | . | 32 | . |
| 08-087 | FPAH | M | 67 | EUR | *CAV1* | NM_001172896.1 | c.381del | p.(Leu128Serfs*22) | frameshift | no | . | 32 | . |
| 11-024 | APAH-CTD | F | 69 | EUR | *CAV1* | NM_001172896.1 | c.407T>C | p.(Phe136Ser) | D-Mis | no | 4.13E-05 | 27 | 0.90 |
| 13-062 | APAH-Portopulm | M | 59 | EUR | *CAV1* | NM_001172896.1 | c.407T>C | p.(Phe136Ser) | D-Mis | no | 4.13E-05 | 27 | 0.90 |
| 16-008 | IPAH | F | 45 | AFR | *CAV1* | NM_001172896.1 | c.418C>T | p.(Arg140Cys) | D-Mis | no | 8.27E-06 | 33 | 0.82 |
| 12-014** | APAH-CTD | F | 78 | EUR | *EIF2AK4* | NM_001013703.3 | c.220G>A | p.(Asp74Asn) | D-Mis | no | 1.66E-05 | 32 | 0.17 |
| 12-014** | APAH-CTD | F | 78 | EUR | *EIF2AK4* | NM_001013703.3 | c.3111G>C | p.(Gln1037His) | D-Mis | no | 8.33E-06 | 25 | 0.51 |
| 02-030** | APAH-CTD | F | 42 | AFR | *EIF2AK4* | NM_001013703.3 | c.650C>G | p.(Pro217Arg) | D-Mis | no | 2.00E-04 | 23 | 0.03 |
| 02-030** | APAH-CTD | F | 42 | AFR | *EIF2AK4* | NM_001013703.3 | c.1265T>G | p.(Val422Gly) | D-Mis | no | 3.00E-04 | 29 | 0.92 |
| 10-091** | FPAH-PVOD | M | 36 | EUR | *EIF2AK4* | NM_001013703.3 | c.1153dup | p.(Val385Glyfs*30) | frameshift | [43] | 3.31E-05 | 32 | . |
| 12-064** | PVOD | F | 48 | EUR | *EIF2AK4* | NM_001013703.3 | c.2141C>A | p.(Ser714*) | stopgain | no | . | 42 | . |
| 10-091** | FPAH-PVOD | M | 36 | EUR | *EIF2AK4* | NM_001013703.3 | c.3766C>T | p.(Arg1256*) | stopgain | [2, 6, 43] | 1.00E-04 | 49 | . |
| 21-036** | IPAH | F | 33 | EUR | *EIF2AK4* | NM_001013703.3 | c.4593del | p.(Ile1533Leufs*2) | frameshift | no | . | 27 | . |
| 12-009 | APAH-HHT | F | 78 | EUR | *ENG* | NM_000118.3 | c.277C>T | p.(Arg93*) | stopgain | [1, 6, 44] | . | 37 | . |
| 06-114 | APAH-HHT | F | 38 | AMR | *ENG* | NM_000118.3 | c.715dup | p.(Glu239Glyfs*95) | frameshift | [44] | . | 28 | . |
| 03-023 | APAH-CTD | F | 69 | EUR | *ENG* | NM_000118.3 | c.1361T>G | p.(Leu454Arg) | D-Mis | no | . | 26 | 0.61 |
| 17-051 | APAH-HIV | M | 57 | AFR | *ENG* | NM_000118.3 | c.1415A>T | p.(Gln472Leu) | D-Mis | no | . | 25 | 0.60 |
| 22-046 | APAH-CTD | F | 53 | EUR | *ENG* | NM_000118.3 | c.1585C>T | p.(Arg529Cys) | D-Mis | no | 1.65E-05 | 29 | 0.64 |
| 38-002 | APAH-CTD | M | 68 | AFR | *ENG* | NM_000118.3 | c.*202_*210dup | p.(=) | splicing | no | 8.03E-05 | 28 | . |
| 25-007 | FPAH | M | 6 | EUR | *KCNK3* | NM_002246.2 | c.544G>A | p.(Glu182Lys) | D-Mis | [6, 45] | . | 32 | 0.59 |
| 15-043 | IPAH | F | 5 | EUR | *KCNK3* | NM_002246.2 | c.646_651dup | p.(Thr216_Gln217dup) | in-frame | [6] | . | 13 | . |
| 05-138 | APAH-CTD | F | 49 | EUR | *KCNK3* | NM_002246.2 | c.1075_1076del | p.(Arg359Thrfs*278) | frameshift | no | . | 35 | . |
| 14-047 | IPAH | M | 62 | EUR | *SMAD4* | NM_005359.5 | c.466_468del | p.(Met157del) | in-frame | no | . | 21 | . |
| 17-009 | IPAH | F | 52 | EUR | *SMAD4* | NM_005359.5 | c.1460C>T | p.(Ala487Val) | D-Mis | no | . | 24 | 0.67 |
| 23-009 | APAH-CTD | F | 45 | EUR | *SMAD9* | NM_001127217.2 | c.138_141del | p.(Lys47Argfs*43) | frameshift | no | 1.65E-05 | 27 | |
| 04-092 | IPAH | M | 58 | AFR | *SMAD9* | NM_001127217.2 | c.146_148del | p.(Lys49_Gly50delinsArg) | in-frame | no | 1.65E-05 | 14 | |
| 06-124 | APAH-CHD | F | 1 | Other | *SMAD9* | NM_001127217.2 | c.204C>A | p.(Cys68*) | stopgain | no | . | 32 | . |
| 11-007 | APAH-HIV | F | 44 | AFR | *SMAD9* | NM_001127217.2 | c.430G>T | p.(Val144Leu) | D-Mis | no | 5.78E-05 | 28 | 0.70 |
| 02-050 | APAH-CTD | F | 47 | AMR | *SMAD9* | NM_001127217.2 | c.438A>C | p.(Arg146Ser) | D-Mis | no | . | 25 | 0.81 |

| ID | PAH subclass | Gender | Age dx | Ancestry | Gene | Transcript | Nucleotide change | Amino acid change | Variant type | Previously reported? | MAF (ExAC) | CADD | REVEL |
|---|---|---|---|---|---|---|---|---|---|---|---|---|---|
| 02-022 | IPAH | F | 57 | AMR | *SMAD9* | NM_001127217.2 | c.804G>C | p.(Glu268Asp) | D-Mis | no | . | 23 | 0.63 |
| 23-004 | APAH-CHD | F | 42 | EUR | *SMAD9* | NM_001127217.2 | c.851G>T | p.(Arg284Leu) | D-Mis | no | 1.65E-05 | 34 | 0.92 |
| 18-070 | APAH-CHD | F | 65 | EAS | SMAD9 | NM_001127217.2 | c.767C>A | p.(Ser256Leu) | D-Mis | no | . | 29 | 0.82 |
| 15-051* | IPAH | F | 4 | EUR | *SMAD9* | NM_001127217.2 | c.880C>T | p.(Arg294*) | stopgain | no | 8.24E-06 | 40 | . |
| 28-062 | APAH-CTD | F | 45 | EUR | *SMAD9* | NM_001127217.2 | c.971C>T | p.(Thr324Met) | D-Mis | no | 8.24E-05 | 28 | 0.79 |
| 26-019 | APAH-CTD | F | 46 | AMR | *SMAD9* | NM_001127217.2 | c.995_997del | p.(Ile332_Gly333delinsArg) | frameshift | no | . | 35 | . |
| 16-040 | IPAH | F | 59 | EUR | *SMAD9* | NM_001127217.2 | c.1260G>C | p.(Lys420Asn) | D-Mis | no | . | 31 | 0.86 |
| 22-006 | FPAH | F | 39 | AMR | *SMAD9* | NM_001127217.2 | c.1321C>T | p.(His441Tyr) | D-Mis | no | . | 29 | 0.92 |
| 15-078 | DTOX | F | 2 | Dominican | *TBX4* | NM_018488.3 | c.146del | p.(Gly49Aspfs*39) | frameshift | no | . | 26 | . |
| 06-061 | IPAH | M | 3 | EUR | *TBX4* | NM_018488.3 | c.150del | p.(Ala52Profs*36) | frameshift | [6] | . | 25 | . |
| 05-184 | DTOX | F | 45 | EUR | *TBX4* | NM_018488.3 | c.179_180dup | p.(Glu61Argfs*28) | frameshift | no | . | 28 | . |
| 02-191 | IPAH | F | 5 | AMR | *TBX4* | NM_018488.3 | c.210dup | p.(Leu71Alafs*3) | frameshift | no | . | 35 | . |
| 08-001 | IPAH | F | 40 | EUR | *TBX4* | NM_018488.3 | c.293C>T | p.(Pro98Leu) | D-Mis | no | . | 34 | 0.97 |
| 02-071* | FPAH | F | 40 | EUR | *TBX4* | NM_018488.3 | c.299A>G | p.(Tyr100Cys) | D-Mis | no | . | 27 | 0.73 |
| 15-013 | IPAH | M | 13 | EUR | *TBX4* | NM_018488.3 | c.316G>A | p.(Gly106Ser) | D-Mis | no | 4.12E-05 | 31 | 0.96 |
| 06-009 | IPAH | F | 14 | AFR | *TBX4* | NM_018488.3 | c.380A>C | p.(Tyr127Ser) | D-Mis | [6] | . | 28 | 0.93 |
| 12-080 | APAH-CTD | F | 29 | AMR | *TBX4* | NM_018488.3 | c.455C>T | p.(Pro152Leu) | D-Mis | no | 1.65E-05 | 34 | 0.95 |
| 10-024 | DTOX | F | 71 | EUR | *TBX4* | NM_018488.3 | c.529C>T | p.(His177Tyr) | D-Mis | no | . | 20 | 0.57 |
| 12-145 | IPAH | M | 70 | EUR | *TBX4* | NM_018488.3 | c.561_570dup | p.(Lys191Leufs*13) | frameshift | no | . | 35 | . |
| 02-159 | APAH-CTD | F | 15 | EUR | *TBX4* | NM_018488.3 | c.571_576del | p.(Lys191_Tyr192del) | in-frame | no | . | 23 | . |
| 15-046 | APAH-CHD | F | 0 | EUR | *TBX4* | NM_018488.3 | c.702+1G>A | p.(=) | splicing | [6, 19] | . | 27 | . |
| 19-038 | APAH-CHD | F | 59 | EUR | *TBX4* | NM_018488.3 | c.782G>A | p.(Arg261Gln) | D-Mis | no | 8.24E-06 | 34 | 0.66 |
| 24-005 | FPAH | M | 15 | Dominican | *TBX4* | NM_018488.3 | c.789del | p.(Ser264Alafs*6) | frameshift | no | . | 33 | . |
| 10-109 | IPAH | F | 57 | AMR | *TBX4* | NM_018488.3 | c.809T>G | p.(Ile270Ser) | D-Mis | no | . | 27 | 0.62 |
| 05-151 | IPAH | M | 63 | EUR | *TBX4* | NM_018488.3 | c.847dup | p.(Gln283Profs*103) | frameshift | [6] | . | 24 | . |
| 15-062 | IPAH | F | 5 | EUR | *TBX4* | NM_018488.3 | c.847dup | p.(Gln283Profs*103) | frameshift | [6] | . | 24 | . |
| 01-008 | IPAH | F | 7 | EUR | *TBX4* | NM_018488.3 | c.985G>T | p.(Asp329Tyr) | D-Mis | no | . | 30 | 0.61 |
| 12-102 | FPAH | M | 55 | EUR | *TBX4* | NM_018488.3 | c.1021+1G>A | p.(=) | splicing | no | . | 27 | . |
| 06-052 | APAH-CHD | M | 1 | EUR | *TBX4* | NM_018488.3 | c.1112del | p.(Pro371Leufs*8) | frameshift | [19] | . | 33 | . |
| 12-045 | IPAH | F | 52 | EUR | *TBX4* | NM_018488.3 | c.1119C>A | p.(Tyr373*) | stopgain | no | . | 36 | . |

| ID | PAH subclass | Gender | Age dx | Ancestry | Gene | Transcript | Nucleotide change | Amino acid change | Variant type | Previously reported? | MAF (ExAC) | CADD | REVEL |
|---|---|---|---|---|---|---|---|---|---|---|---|---|---|
| 02-168 | IPAH | M | 17 | EAS | *TBX4* | NM_018488.3 | c.1417_1420dup | p.(Tyr474Cysfs*30) | frameshift | no | . | 21 | . |

Rare, predicted deleterious variants defined as MAF ≤1.00E-04 and LGD (stopgain, frameshift, splicing) or missense with REVEL score >0.5 (D-Mis). For biallelic *EIF2AK4* variants, MAF ≤1.00E-02 and LGD or missense with CADD ≥20.

*Patients 02-071, 02-100, 12-015 and 15-051 carried variants in more than one risk gene.

**Patients 12-064 and 21-036 are homozygous for the given EIF2AK4 mutations. Patients 12-014, 02-030 and 10-091 are compound heterozygotes.

Abbreviations: dx, diagnosis; HHT, hereditary hemorrhagic telangiectasia; MAF, minor allele frequency; NF, neurofibromatosis

**Supplementary Table 3. Rare, predicted deleterious variants in recently-reported PAH risk genes among 2572 PAH cases. Patients are heterozygous for the indicated variant.**

| Patient ID | PAH subclass | Gender | Age at dx | Ancestry | Gene | Transcript | Nucleotide change | Amino acid change | Variant type | Previously reported? | MAF (ExAC) | CADD | REVEL |
|---|---|---|---|---|---|---|---|---|---|---|---|---|---|
| 18-002 | IPAH | F | 34 | EAS | *ABCC8* | NM_001351295.1 | c.307C>T | p.(His103Tyr) | D-Mis | rs751209734 | 4.95E-05 | 23 | 0.66 |
| 05-145 | APAH-CHD | F | 42 | AMR | *ABCC8* | NM_001351295.1 | c.313C>T | p.(His105Tyr) | D-Mis | no | 1.65E-05 | 27 | 0.78 |
| 12-207 | IPAH | F | 43 | EUR | *ABCC8* | NM_001351295.1 | c.375C>G | p.(His125Gln) | D-Mis | rs60637558 | 8.25E-05 | 26 | 0.89 |
| 05-166 | APAH-CTD | F | 31 | SAS | *ABCC8* | NM_001351295.1 | c.503G>A | p.(Arg168His) | D-Mis | no | . | 30 | 0.88 |
| 02-060 | IPAH | F | 34 | AMR | *ABCC8* | NM_001351295.1 | c.610G>T | p.(Val204Leu) | D-Mis | no | . | 20 | 0.54 |
| 22-023 | IPAH | F | 33 | EUR | *ABCC8* | NM_001351295.1 | c.647G>A | p.(Arg216His) | D-Mis | rs199702708 | 2.47E-05 | 24 | 0.60 |
| 08-043 | IPAH | M | 57 | EUR | *ABCC8* | NM_001351295.1 | c.890G>T | p.(Arg297Met) | D-Mis | no | 8.30E-06 | 26 | 0.82 |
| 13-078 | APAH-CTD | F | 41 | AFR | *ABCC8* | NM_001351295.1 | c.1024G>A | p.(Gly342Arg) | D-Mis | no | 3.31E-05 | 23 | 0.63 |
| 02-179 | APAH-CTD | F | 50 | EUR | *ABCC8* | NM_001351295.1 | c.1198A>G | p.(Met400Val) | D-Mis | no | . | 22 | 0.55 |
| 05-033 | IPAH | F | 63 | EUR | *ABCC8* | NM_001351295.1 | c.1198A>G | p.(Met400Val) | D-Mis | no | . | 22 | 0.55 |
| 07-025 | APAH-CTD | F | 64 | AFR | *ABCC8* | NM_001351295.1 | c.1270G>A | p.(Asp424Asn) | D-Mis | rs577545383 | 2.47E-05 | 34 | 0.96 |
| 13-034 | IPAH | F | 73 | EUR | *ABCC8* | NM_001351295.1 | c.1484G>A | p.(Arg495Gln) | D-Mis | no | . | 35 | 0.87 |
| 14-006 | APAH-HIV | M | 44 | EUR | *ABCC8* | NM_001351295.1 | c.1531C>G | p.(Leu511Val) | D-Mis | no | 8.25E-06 | 27 | 0.82 |
| 06-043 | APAH-CHD | F | 44 | AMR | *ABCC8* | NM_001351295.1 | c.1676T>C | p.(Phe559Ser) | D-Mis | no | . | 32 | 0.95 |
| 16-054 | IPAH | F | 44 | AFR | *ABCC8* | NM_001351295.1 | c.1919C>A | p.(Ala640Glu) | D-Mis | no | 8.26E-06 | 16 | 0.54 |
| 17-018 | APAH-CTD | F | 48 | EUR | *ABCC8* | NM_001351295.1 | c.2003T>C | p.(Val668Ala) | D-Mis | no | . | 23 | 0.57 |
| 18-025 | IPAH | F | 33 | EAS | *ABCC8* | NM_001351295.1 | c.2008C>T | p.(Arg670Cys) | D-Mis | no | 8.31E-06 | 34 | 0.73 |
| 19-051 | IPAH | F | 29 | AMR | *ABCC8* | NM_001351295.1 | c.2008C>T | p.(Arg670Cys) | D-Mis | no | 8.31E-06 | 34 | 0.73 |
| 11-009 | APAH-CTD | F | 76 | EUR | *ABCC8* | NM_001351295.1 | c.2218G>A | p.(Gly740Ser) | D-Mis | no | 1.66E-05 | 35 | 0.96 |
| 16-036 | APAH-CTD | F | 64 | EUR | *ABCC8* | NM_001351295.1 | c.2228C>T | p.(Ser743Leu) | D-Mis | no | . | 34 | 0.95 |
| 06-018 | IPAH | M | 18 | EUR | *ABCC8* | NM_001351295.1 | c.2230C>T | p.(Leu744Phe) | D-Mis | no | 8.26E-06 | 29 | 0.90 |
| 31-006 | APAH-CTD | F | 50 | AFR | *ABCC8* | NM_001351295.1 | c.2439T>A | p.(Ser813Arg) | D-Mis | no | . | 17 | 0.59 |
| 04-056 | IPAH | F | 41 | EUR | *ABCC8* | NM_001351295.1 | c.2474A>G | p.(Glu825Gly) | D-Mis | no | . | 23 | 0.56 |
| 15-025 | APAH-CHD | F | 2 | EUR | *ABCC8* | NM_001351295.1 | c.2488C>A | p.(Gln830Lys) | D-Mis | no | 4.12E-05 | 20 | 0.60 |
| 07-055 | IPAH | F | 67 | EUR | *ABCC8* | NM_001351295.1 | c.2488C>A | p.(Gln830Lys) | D-Mis | no | 4.12E-05 | 20 | 0.60 |
| 06-091 | APAH-CHD | F | 15 | AFR | *ABCC8* | NM_001351295.1 | c.3410C>T | p.(Thr1137Met) | D-Mis | no | 8.25E-06 | 34 | 0.61 |
| 03-076 | APAH-CTD | F | 64 | EUR | *ABCC8* | NM_001351295.1 | c.3571C>A | p.(Pro1191Thr) | D-Mis | no | . | 27 | 0.84 |
| 20-035 | IPAH | F | 35 | AFR | *ABCC8* | NM_001351295.1 | c.3644A>T | p.(Asp1215Val) | D-Mis | no | 2.47E-05 | 29 | 0.94 |
| 06-052 | IPAH | M | 1 | EUR | *ABCC8* | NM_001351295.1 | c.4004G>A | p.(Arg1335His) | D-Mis | [46] | 2.48E-05 | 34 | 0.93 |
| 12-063 | APAH-CTD | F | 30 | EUR | *ATP13A3* | NM_024524.3 | c.158_159del; 163_167del | p.(Trp53Serfs*12) | frameshift | no | . | 27 | . |

| Patient ID | PAH subclass | Gender | Age at dx | Ancestry | Gene | Transcript | Nucleotide change | Amino acid change | Variant type | Previously reported? | MAF (ExAC) | CADD | REVEL |
|---|---|---|---|---|---|---|---|---|---|---|---|---|---|
| 07-091 | IPAH | F | 40 | EUR | *ATP13A3* | NM_024524.3 | c.201_202del | p.(Cys67*) | stopgain | no | . | 25 | . |
| 03-012 | APAH-CHD | M | 30 | EUR | *ATP13A3* | NM_024524.3 | c.1222A>G | p.(Arg408Gly) | D-Mis | no | . | 26 | 0.72 |
| 04-025 | IPAH | F | 66 | EUR | *ATP13A3* | NM_024524.3 | c.2189_2205del | p.(Thr730Argfs*4) | frameshift | no | . | 35 | . |
| 14-040 | IPAH | F | 56 | SAS | *ATP13A3* | NM_024524.3 | c.2228G>T | p.(Arg743Leu) | D-Mis | no | . | 35 | 0.75 |
| 09-043 | IPAH | F | 45 | EUR | *ATP13A3* | NM_024524.3 | c.2549dup | p.(Met850Ilefs*13) | frameshift | no | . | 35 | . |
| 06-014 | FPAH | M | 35 | EUR | *ATP13A3* | NM_024524.3 | c.2996C>T | p.(Ser999Leu) | D-Mis | no | 8.30E-06 | 29 | 0.74 |
| 18-055 | IPAH | M | 53 | AMR | *GDF2* | NM_016204.3 | c.76C>T | p.(Gln26*) | stopgain | [47] | . | 36 | . |
| 26-003 | IPAH | F | 39 | EUR | *GDF2* | NM_016204.3 | c.143dup | p.(Pro49Alafs*2) | frameshift | no | . | 23 | . |
| 18-029 | IPAH | F | 22 | SAS | *GDF2* | NM_016204.3 | c.230C>T | p.(Ser77Leu) | D-Mis | no | . | 22 | . |
| 34-016 | APAH-CTD | F | 67 | AMR | *GDF2* | NM_016204.3 | c.254C>T | p.(Pro85Leu) | D-Mis | no | . | 24 | . |
| 02-116 | IPAH | F | 55 | EUR | *GDF2* | NM_016204.3 | c.328C>T | p.(Arg110Trp) | D-Mis | [8] | . | 31 | . |
| 05-003 | IPAH | M | 48 | EUR | *GDF2* | NM_016204.3 | c.329G>A | p.(Arg110Gln) | D-Mis | no | . | 34 | . |
| 10-056 | IPAH | M | 26 | AMR | *GDF2* | NM_016204.3 | c.502G>T | p.(Gly168*) | stopgain | no | . | 37 | . |
| 28-107 | IPAH | F | 39 | AMR | *GDF2* | NM_016204.3 | c.530A>T | p.(Asp177Val) | D-Mis | no | . | 26 | . |
| 05-109 | IPAH | F | 22 | AMR | *GDF2* | NM_016204.3 | c.607G>C | p.(Glu203Gln) | D-Mis | no | . | 24 | . |
| 24-022 | IPAH | M | 9 | EAS | *GDF2* | NM_016204.3 | c.646C>T | p.(Arg216Trp) | D-Mis | no | . | 24 | . |
| 13-058 | APAH-HIV | M | 70 | EUR | *GDF2* | NM_016204.3 | c.751del | p.(Leu251Cysfs*22) | frameshift | no | . | 24 | . |
| 12-083 | IPAH | F | 46 | EUR | *GDF2* | NM_016204.3 | c.751del | p.(Leu251Cysfs*22) | frameshift | no | . | 24 | . |
| 03-122 | IPAH | F | 45 | EUR | *GDF2* | NM_016204.3 | c.857dup | p.(Leu287Alafs*11) | frameshift | no | . | 24 | . |
| 27-008 | IPAH | F | 41 | EUR | *GDF2* | NM_016204.3 | c.857dup | p.(Leu287Alafs*11) | frameshift | no | . | 24 | . |
| 31-021 | APAH-HIV | F | 52 | AFR | *GDF2* | NM_016204.3 | c.997C>T | p.(Arg333Trp) | D-Mis | no | 1.65E-05 | 22 | . |
| 18-079 | IPAH | F | 31 | AMR | *GDF2* | NM_016204.3 | c.997C>T | p.(Arg333Trp) | D-Mis | no | 1.65E-05 | 22 | . |
| 02-039 | IPAH | F | 45 | AMR | *GDF2* | NM_016204.3 | c.1011G>T | p.(Glu337Asp) | D-Mis | no | . | 23 | . |
| 02-039 | IPAH | F | 45 | AMR | *GDF2* | NM_016204.3 | c.1012G>T | p.(Asp338Tyr) | D-Mis | no | . | 28 | . |
| 29-045 | FPAH | F | 64 | EUR | *GDF2* | NM_016204.3 | c.1023G>C | p.(Trp341Cys) | D-Mis | no | . | 35 | . |
| 06-014 | FPAH | M | 35 | EUR | *GDF2* | NM_016204.3 | c.1042C>A | p.(Pro348Thr) | D-Mis | no | . | 25 | . |
| 27-002 | IPAH | M | 59 | EUR | *GDF2* | NM_016204.3 | c.1063G>C | p.(Glu355Gln) | D-Mis | no | . | 25 | . |
| 09-073 | IPAH | F | 18 | AFR | *GDF2* | NM_016204.3 | c.1103T>C | p.(Val368Ala) | D-Mis | no | . | 24 | . |
| 02-173 | APAH-CTD | F | 51 | AFR | *GDF2* | NM_016204.3 | c.1135del | p.(Leu379fs) | frameshift | no | . | . | . |
| 17-089 | APAH-HIV | F | 43 | AFR | *GDF2* | NM_016204.3 | c.1135del | p.(Leu379fs) | frameshift | no | . | . | . |
| 17-017 | IPAH | F | 33 | EUR | *GDF2* | NM_016204.3 | c.1186A>G | p.(Thr396Ala) | D-Mis | no | . | 29 | . |

| Patient ID | PAH subclass | Gender | Age at dx | Ancestry | Gene | Transcript | Nucleotide change | Amino acid change | Variant type | Previously reported? | MAF (ExAC) | CADD | REVEL |
|---|---|---|---|---|---|---|---|---|---|---|---|---|---|
| 05-156 | IPAH | F | 40 | EAS | *GDF2* | NM_016204.3 | c.1259G>C | p.(Gly420Ala) | D-Mis | no | . | 27 | . |
| 09-046 | IPAH | F | 24 | EUR | *GDF2* | NM_016204.3 | c.1267G>A | p.(Val423Met) | D-Mis | no | . | 29 | . |
| 02-129 | IPAH | F | 25 | EUR | *GDF2* | NM_016204.3 | c.1282T>C | p.(Cys428Arg) | D-Mis | no | . | 31 | . |
| 05-174 | APAH-CTD | F | 56 | AMR | *KCNA5* | NM_002234.3 | c.1A>C | (p.?) | D-Mis | no | . | 23 | 0.55 |
| 28-020 | IPAH | F | 21 | EUR | *KCNA5* | NM_002234.3 | c.660T>G | p.(Ile220Met) | D-Mis | no | . | 16 | 0.5 |
| 12-050 | FPAH | F | 36 | EUR | *KCNA5* | NM_002234.3 | c.670del | p.(Glu224Argfs*134) | frameshift | no | . | 21 | . |
| 12-130 | FPAH | F | 28 | EUR | *KCNA5* | NM_002234.3 | c.670del | p.(Glu224Argfs*134) | frameshift | no | . | 21 | . |
| 12-140 | FPAH | F | 59 | EUR | *KCNA5* | NM_002234.3 | c.670del | p.(Glu224Argfs*134) | frameshift | no | . | 21 | . |
| 11-060 | APAH-CTD | M | 62 | EUR | *KCNA5* | NM_002234.3 | c.964G>C | p.(Asp322His) | D-Mis | no | 9.15E-05 | 24 | 0.83 |
| 20-028 | APAH-CHD | F | 26 | EUR | *KCNA5* | NM_002234.3 | c.964G>C | p.(Asp322His) | D-Mis | no | 9.15E-05 | 24 | 0.83 |
| 17-098 | APAH-APAH | F | 73 | AFR | *KCNA5* | NM_002234.3 | c.1043G>A | p.(Ser348Asn) | D-Mis | no | 8.25E-06 | 25 | 0.56 |
| 04-081 | APAH-CTD | F | 43 | EUR | *KCNA5* | NM_002234.3 | c.1243C>T | p.(Arg415Cys) | D-Mis | no | . | 32 | 0.84 |
| 21-048 | IPAH | F | 75 | EUR | *KCNA5* | NM_002234.3 | c.1327A>G | p.(Ile443Val) | D-Mis | no | 5.78E-05 | 25 | 0.67 |
| 05-179 | APAH-CTD | F | 50 | EUR | *KCNA5* | NM_002234.3 | c.1472T>C | p.(Val491Ala) | D-Mis | no | 4.94E-05 | 23 | 0.55 |
| 18-014 | DTOX | M | 45 | EUR | *KCNA5* | NM_002234.3 | c.1564T>C | p.(Phe522Leu) | D-Mis | no | . | 29 | 0.93 |
| 12-017 | APAH-CHD | F | 30 | EAS | *KCNA5* | NM_002234.3 | c.1727C>T | p.(Ala576Val) | D-Mis | no | 3.33E-05 | 13 | 0.54 |
| 07-020 | APAH-CHD | F | 53 | EUR | *SMAD1* | NM_005900.2 | c.469C>T | p.(Arg157Cys) | D-Mis | no | 8.25E-06 | 32 | 0.56 |
| 06-089 | APAH-CTD | F | 50 | EUR | *SMAD1* | NM_005900.2 | c.671C>A | p.(Pro224Gln) | D-Mis | no | . | 24 | 0.71 |
| 06-116 | APAH-CHD | F | 3 | EUR | *SOX17* | NM_022454.3 | c.226A>G | p.(Met76Val) | D-Mis | [19] | . | 26 | 0.97 |
| 29-021 | APAH-Portopulm | F | 57 | AMR | *SOX17* | NM_022454.3 | c.277C>A | p.(Leu93Met) | D-Mis | no | . | 26 | 0.63 |
| 24-002 | IPAH | F | 7 | EUR | *SOX17* | NM_022454.3 | c.365_366del | p.(Glu122Alafs*39) | frameshift | no | . | 34 | . |
| 28-150 | APAH-CHD | F | 31 | AMR | *SOX17* | NM_022454.3 | c.392A>G | p.(Asp131Gly) | D-Mis | [19] | . | 29 | 0.89 |
| 05-192 | IPAH | F | 40 | AMR | *SOX17* | NM_022454.3 | c.392A>G | p.(Asp131Gly) | D-Mis | [19] | . | 29 | 0.89 |
| 06-005 | IPAH | F | 5 | EUR | *SOX17* | NM_022454.3 | c.418C>T | p.(Arg140Trp) | D-Mis | no | . | 35 | 0.68 |
| 06-012 | IPAH | F | 5 | AMR | *SOX17* | NM_022454.3 | c.499_520del | p.(Leu167Trpfs*213) | frameshift | [8] | . | 34 | . |
| 08-008 | DTOX | M | 31 | EUR | *SOX17* | NM_022454.3 | c.788del | p.(Pro263Argfs*124) | frameshift | no | . | 10 | . |
| 07-059 | IPAH | F | 66 | EUR | *SOX17* | NM_022454.3 | c.1190C>T | p.(Ser397Leu) | D-Mis | no | . | 34 | 0.80 |
| 06-016 | IPAH | M | 19 | EUR | *SOX17* | NM_022454.3 | c.1224del | p.(Cys409Alafs*45) | frameshift | no | . | 33 | . |

Rare, predicted deleterious variants defined as MAF ≤1.00E-04 and LGD (stopgain, frameshift, splicing) or missense with REVEL score >0.5 (D-Mis).
Abbreviations: dx, diagnosis; MAF, minor allele frequency.

Supplementary Table 2 and 3 References

**Supplementary Table 4. Similar frequency of rare synonymous variants among European PAH cases and non-Finnish European gnomAD and in-house controls.**

| Mutation type* | PAH cases (n=1,832) | controls (n=12,771) | Enrichment | p-value |
|---|---|---|---|---|
| SYN | 60024 | 420379 | 1.0 | 0.09 |
| LGD | 8198 | 57417 | 1.0 | 0.53 |
| MIS | 122405 | 862011 | 1.0 | 0.52 |

*Syn, synonymous; LGD, likely gene damaging; Mis, missense.

**Supplementary Table 5A. Enrichment of rare deleterious variants in a *KLK* gene-set* expressed in lung among 1,832 European PAH cases and 12,771 European controls.**

| Mutation type** | PAH cases (n=1,832) | Controls (n=12,771) | Enrichment | p-value |
|---|---|---|---|---|
| LGD | 5 | 12 | 2.9 | 0.05 |
| D-Mis | 16 | 56 | 2.0 | 0.02 |
| LGD+D-Mis | 21 | 69 | 2.1 | 0.004 |

*KLK gene-set: KLK1, KLK5, KLK6, KLK7, KLK10, KLK11, KLK12, KLK13, KLK14.
**LGD, likely gene damaging; D-Mis, missense with REVEL variable threshold.

**Supplementary Table 5B. Association analysis of *KLK* genes expressed in lung using 1,832 (all PAH) or 812 (IPAH) European cases and 12,771 European controls.**

| Gene | PAH subclass | Empirical Revel | P-value | Number of permutations | Permutation p-value | OR |
|---|---|---|---|---|---|---|
| *KLK1* | all PAH | 0.45 | 1.35E-07 | 10000000 | 2.00E-07 | 13.9 |
| *KLK12* | all PAH | 0.7 | 0.03 | 1000 | 0.07 | 7.0 |
| *KLK10* | all PAH | 0.25 | 0.04 | 1000 | 0.09 | 2.4 |
| *KLK13* | all PAH | 0.25 | 0.10 | 1000 | 0.13 | 2.3 |
| *KLK5* | all PAH | 0.4 | 0.33 | 1000 | 0.44 | 1.6 |
| *KLK14* | all PAH | 0.4 | 0.31 | 1000 | 0.46 | 2.0 |
| *KLK6* | all PAH | 0.85 | 0.42 | 1000 | 0.55 | 2.3 |
| *KLK7* | all PAH | 0.2 | 0.35 | 1000 | 0.59 | 1.3 |
| *KLK11* | all PAH | 0.25 | 0.82 | 1000 | 0.89 | 0.6 |
| *KLK1* | IPAH | 0.45 | 3.34E-09 | 10000000 | 1.00E-07 | 26.2 |
| *KLK12* | IPAH | 0.7 | 0.004 | 1000 | 0.007 | 15.7 |
| *KLK13* | IPAH | 0.3 | 0.024 | 1000 | 0.032 | 4.2 |
| *KLK10* | IPAH | 0.25 | 0.052 | 1000 | 0.076 | 3.1 |
| *KLK5* | IPAH | 0.4 | 0.225 | 1000 | 0.339 | 2.4 |
| *KLK14* | IPAH | 0.6 | 0.309 | 1000 | 0.412 | 3.1 |
| *KLK11* | IPAH | 0.25 | 0.758 | 1000 | 0.742 | 0.7 |
| *KLK7* | IPAH | 0.25 | 0.772 | 1000 | 0.767 | 0.7 |
| *KLK6* | IPAH | 0.8 | 1 | 1000 | 1 | 2.0 |

**Supplemental Table 6. Mean clinical phenotypes of *KLK1* and *GGCX* IPAH cases compared to other IPAH cases without variants in known risk genes*.**

| Group | Age dx (y) | MPAP (mmHg) | MPCW (mmHg) | CO, Fick (L/min) | PVR (Woods units) | MAP (mmHg) | MAP:MPAP |
|---|---|---|---|---|---|---|---|
| ***KLK1*** | 48 ± 19 | 46 ± 12 | 10 ± 3 | 4.7 ± 1.7 | 8.2 ± 4.7 | 99 ± 7 | 2.4 ± 0.4 |
| **(n)** | (10) | (10) | (10) | (8) | (8) | (7) | (7) |
| ***GGCX*** | 48 ± 15 | 57 ± 14 | 11 ± 4 | 4.5 ± 1.4 | 10.7 ± 8.0 | 88 ± 17** | 1.6 ± 0.6 |
| **(n)** | (17) | (17) | (15) | (12) | (10) | (10) | (10) |
| **other IPAH** | 48 ± 19 | 52 ± 12 | 12 ± 5 | 7.0 ± 3.2 | 11.3 ± 7.2 | 96 ± 15 | 1.9 ± 0.7 |
| **(n)** | (1096) | (1043) | (1016) | (715) | (731) | (582) | (582) |

Abbreviations: Age dx, patient age at diagnosis; MPAP, mean pulmonary arterial pressure; MPCW, mean pulmonary capillary wedge pressure; CO, cardiac output by Frick method; PVR, pulmonary vascular resistance; MAP, mean arterial pressure.
*known risk genes included 18 genes from Supplementary Tables 2 and 3.
**p<0.02 vs other IPAH, Student's t-test.

**Supplementary Table 7. Lack of enrichment of *KLK1* common SNP, R77H, in the PAH Biobank cohort compared to gnomAD population data.**

| Ancestry group | # R77H alleles observed in cases | # Total cases | # R77H alleles in gnomad | p-value | RR |
|---|---|---|---|---|---|
| NFE | 136 | 1,841 | 0.0371 | 1.00 | 1.0 |
| AFR | 54 | 291 | 0.1033 | 0.54 | 0.9 |

NFE, non-Finnish European; AFR, African; AF, allele frequency; RR, relative risk.
P-value based on binomial test.

**Supplementary Figure Legends**

**Supplementary Figure 1. Locations of rare deleterious PAH patient-derived *BMPR2* variants within the two-dimensional protein structure.** Predicted damaging missense (D-Mis) variants are shown above the protein schematic; likely-gene-disrupting (LGD including stopgain, frameshift, in-frame deletion and whole exon deletion) variants are below the schematic. The vertical gray lines indicate exon borders. E1(2) indicates a deletion of exon 1 identified in two cases; similar designations are used for the other whole exon deletions.

**Supplementary Figure 2. Locations of rare deleterious PAH patient-derived other previously reported PAH risk gene variants within the two-dimensional protein structures.** Predicted damaging missense (D-Mis) variants are shown above the protein schematics; likely-gene-disrupting (LGD including stopgain, frameshift, in-frame deletion and whole exon deletion) variants are shown below the schematics. The vertical gray lines indicate exon borders. For *ACVRL1*, E10 (1) indicates a deletion of exon 10 identified in one case.

**Supplementary Figure 3. Gene-level burden test for rare synonymous variants using 1832 European cases and 12,771 European controls.** Results of a binomial test confined to rare synonymous variants in 20,000 protein-coding genes.

**Supplementary Figure 4. Depth of coding sequence coverage for *GGCX* and *KLK1*.** Comparison of coverage between PAH Biobank cases and gnomAD WGS controls at read depth >10X or >15X.

**Supplementary Figure 1**

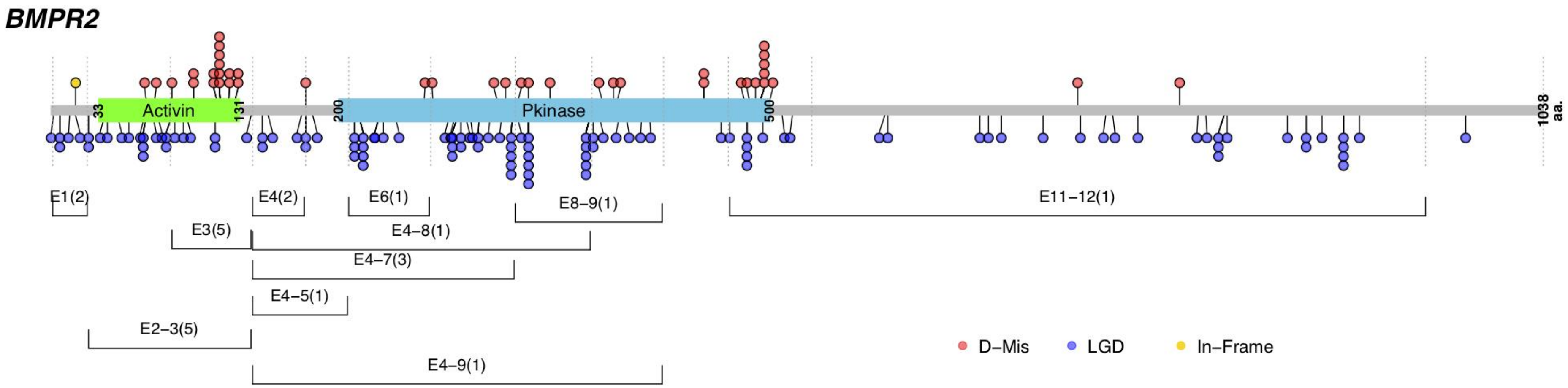

**Supplementary Figure 2**

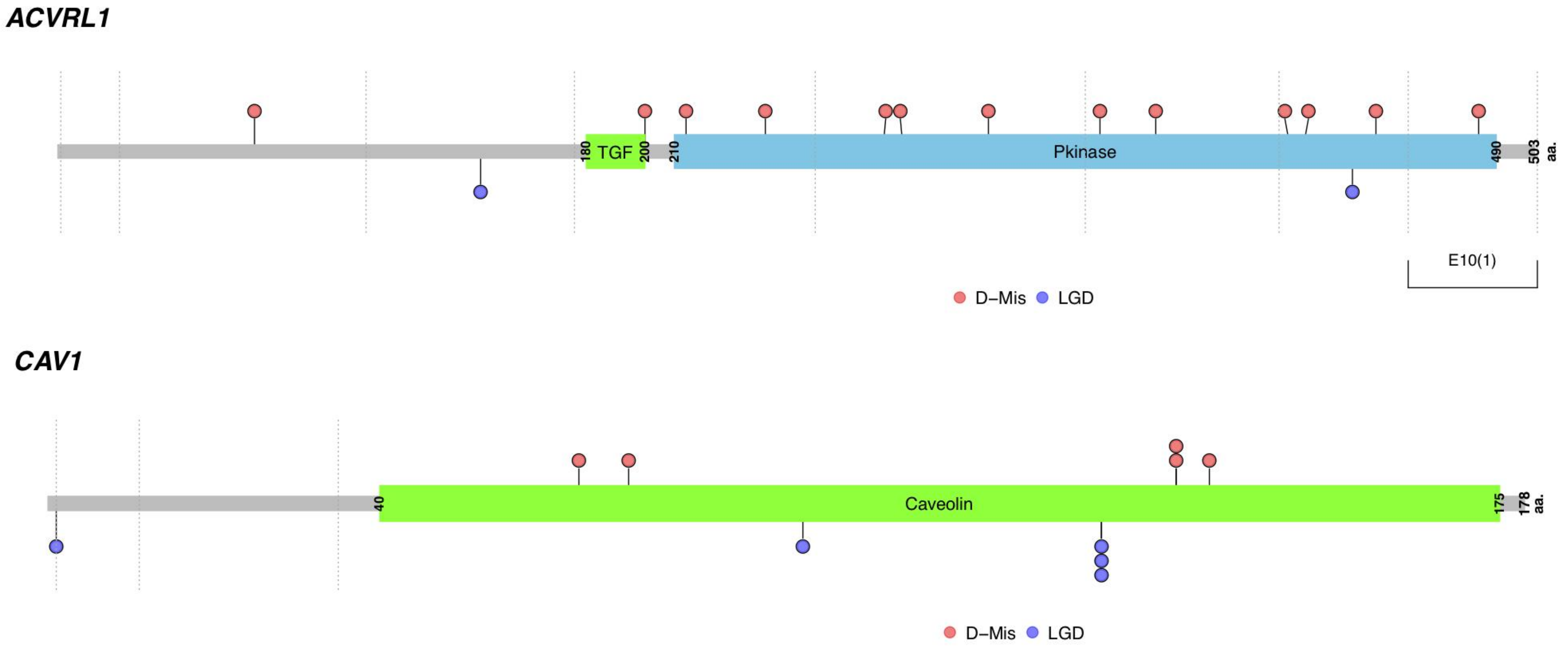

*BMPR1A*

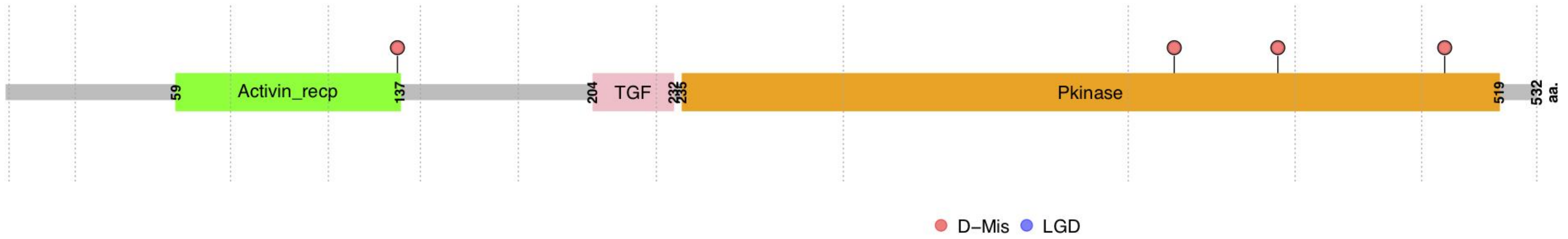

*BMPR1B*

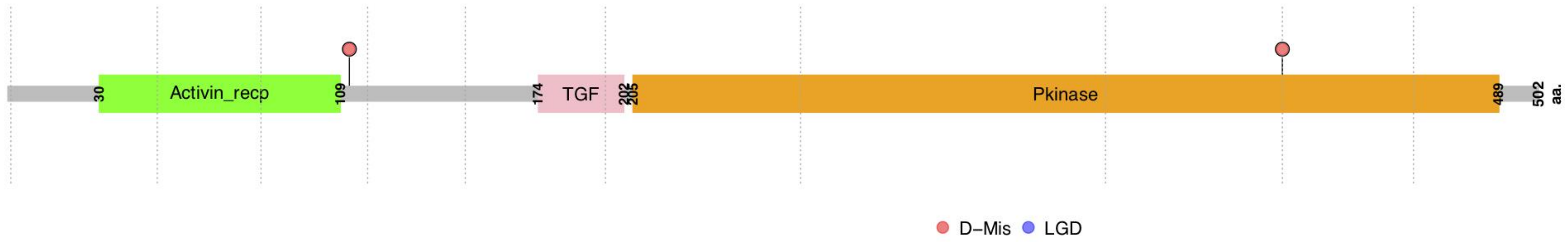

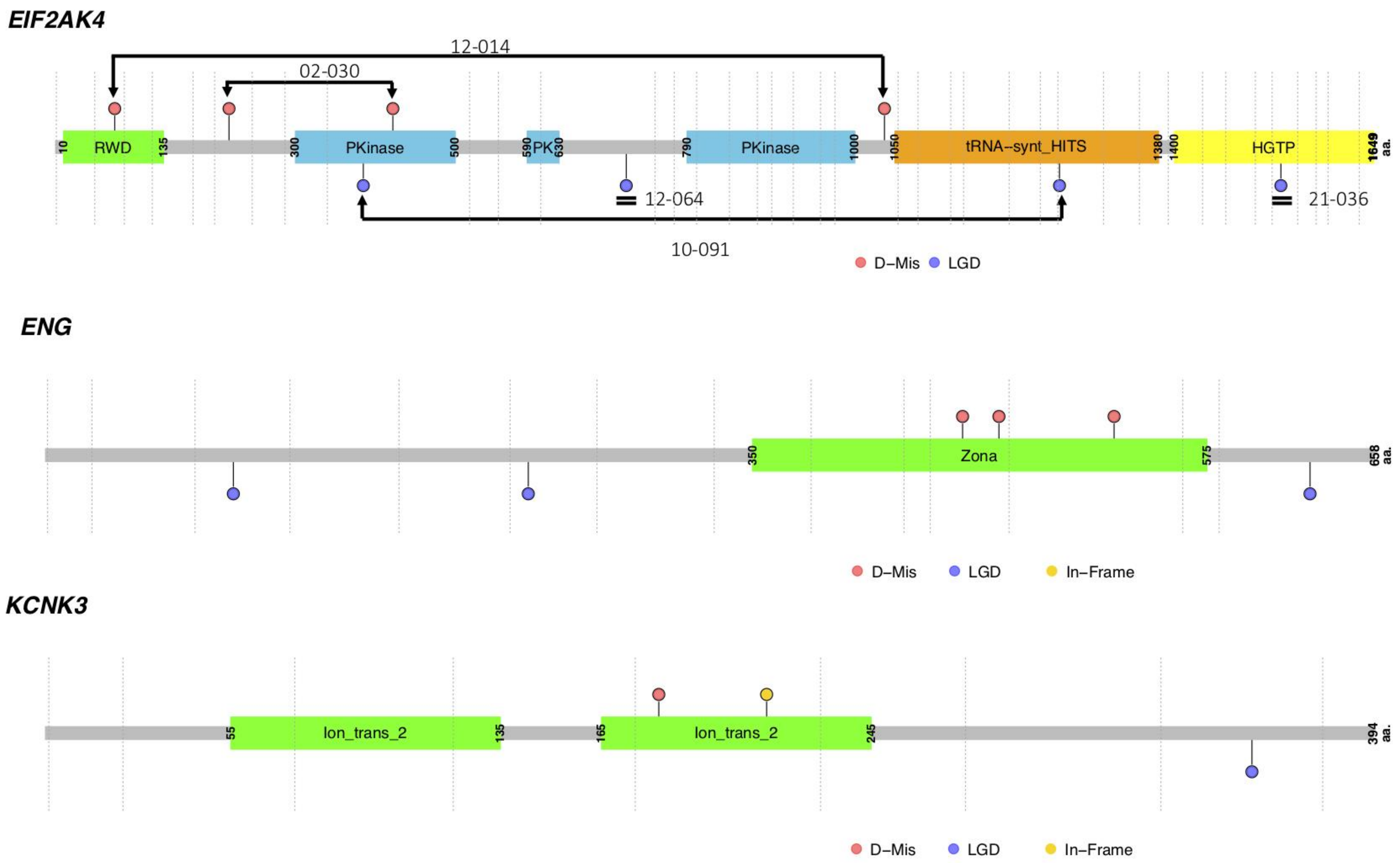
EIF2AK4
12-014
02-030
10
RWD
135
300
PKinase
500
590
PK
630
790
PKinase
1000
1050
tRNA–synt_HITS
1380
1400
HGTP
1649
aa.
12-064
21-036
10-091
D–Mis
LGD
ENG
350
Zona
575
658
aa.
D–Mis
LGD
In–Frame
KCNK3
55
Ion_trans_2
135
165
Ion_trans_2
245
394
aa.
D–Mis
LGD
In–Frame

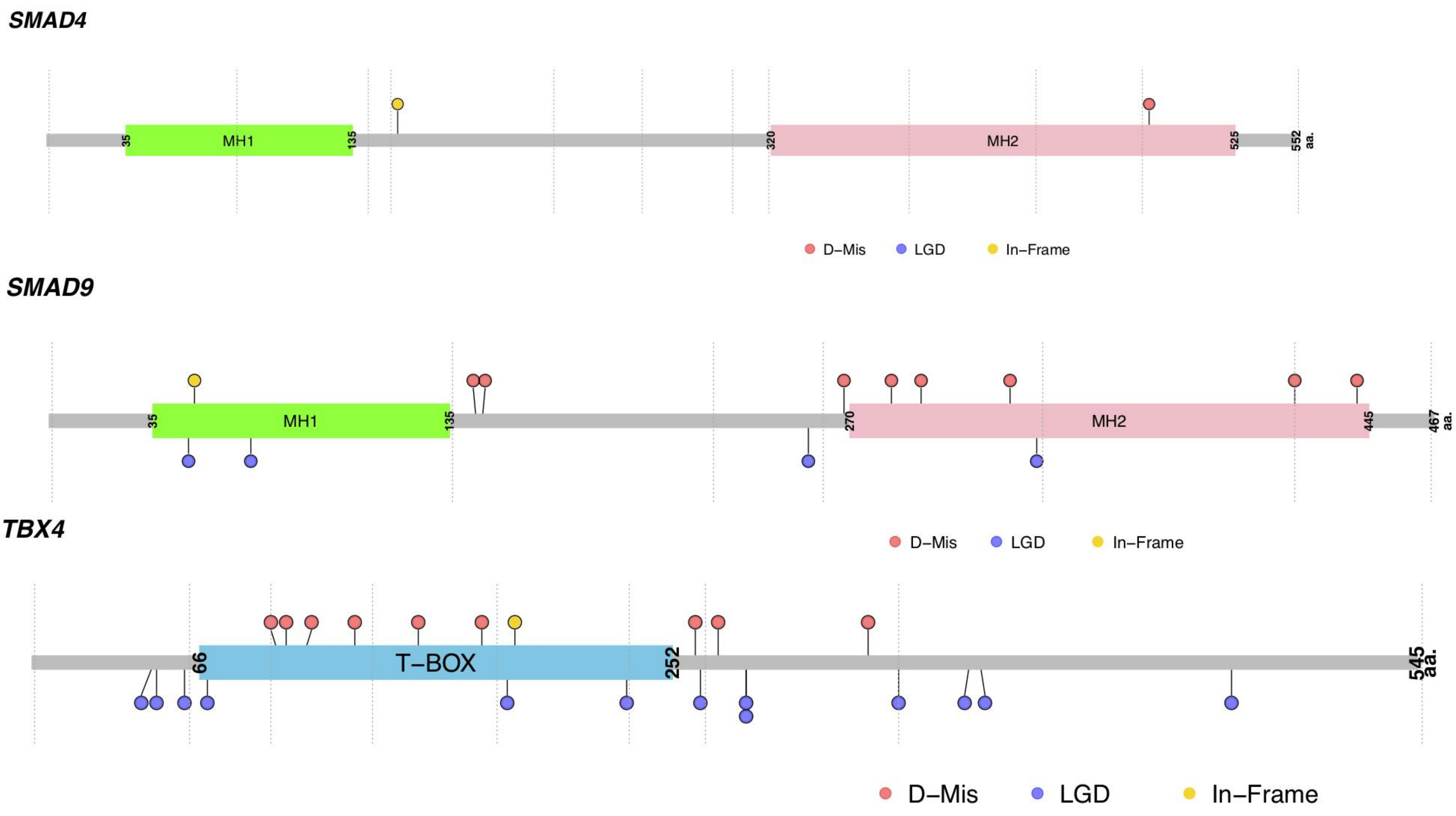
SMAD4
MH1
35
135
MH2
320
525
552 aa.
D−Mis
LGD
In−Frame
SMAD9
MH1
35
135
MH2
270
445
467 aa.
D−Mis
LGD
In−Frame
TBX4
T−BOX
66
252
545 aa.
D−Mis
LGD
In−Frame

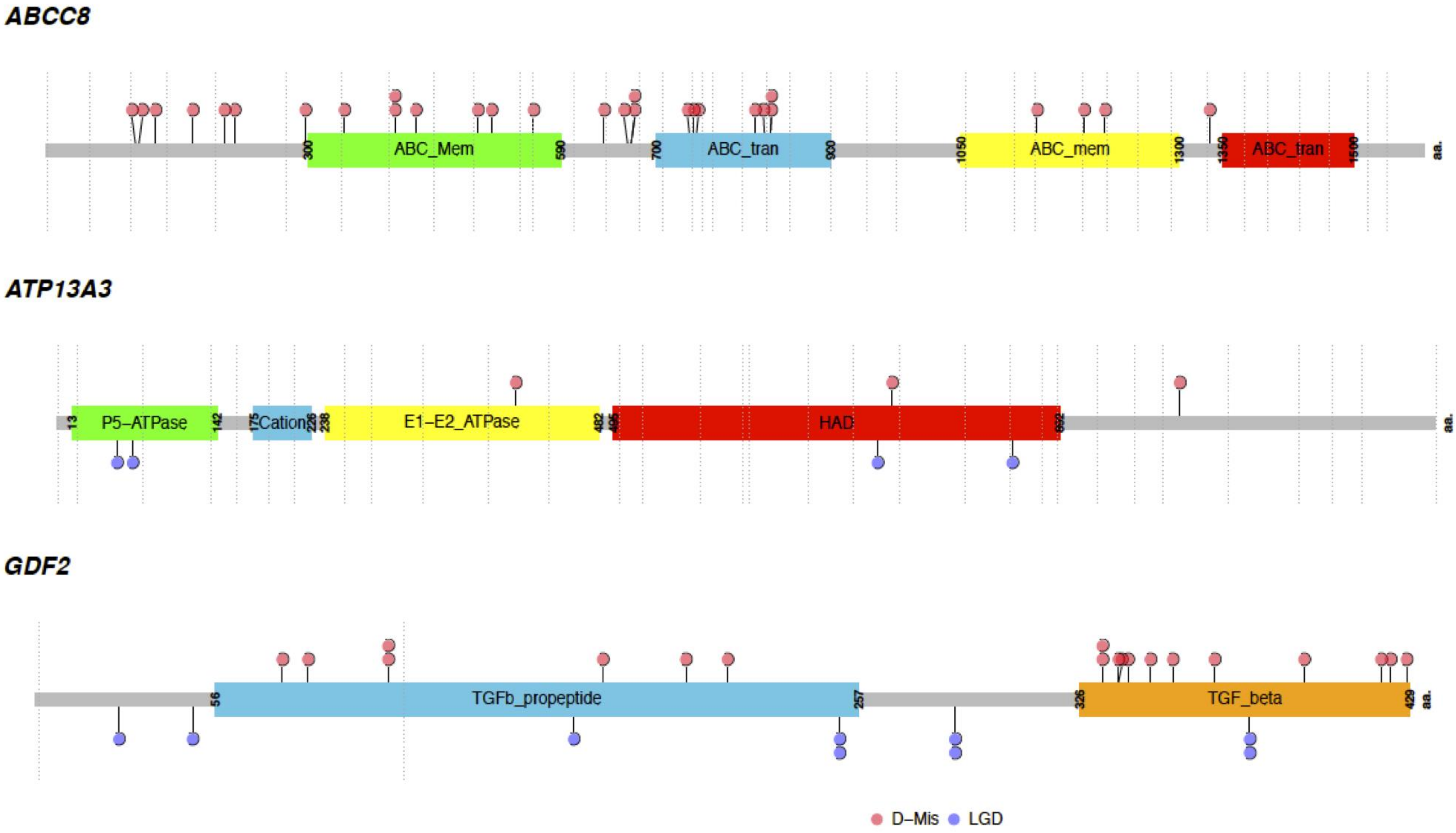

ABCC8
ABC_Mem
ABC_tran
ABC_mem
ABC_tran
300
590
700
900
1050
1300
1350
1500
aa.
ATP13A3
P5–ATPase
Cation
E1–E2_ATPase
HAD
13
142
175
226
238
482
aa.
GDF2
TGFb_propeptide
TGF_beta
56
257
326
429
aa.
D–Mis
LGD

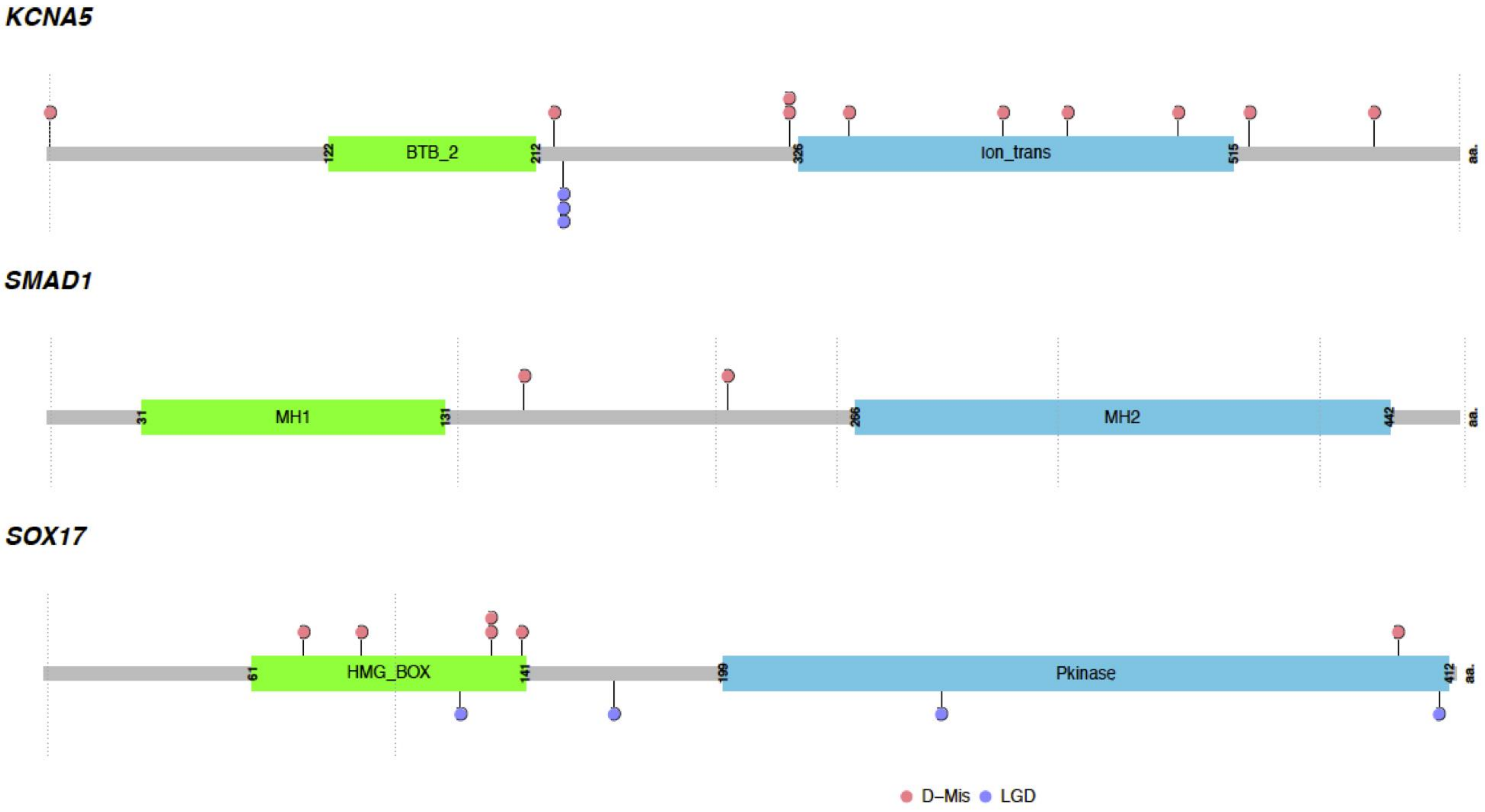
KCNA5
122
BTB_2
212
326
Ion_trans
515
aa.
SMAD1
31
MH1
131
266
MH2
442
aa.
SOX17
61
HMG_BOX
141
199
Pkinase
412
aa.
D–Mis
LGD

**Supplementary Figure 3**

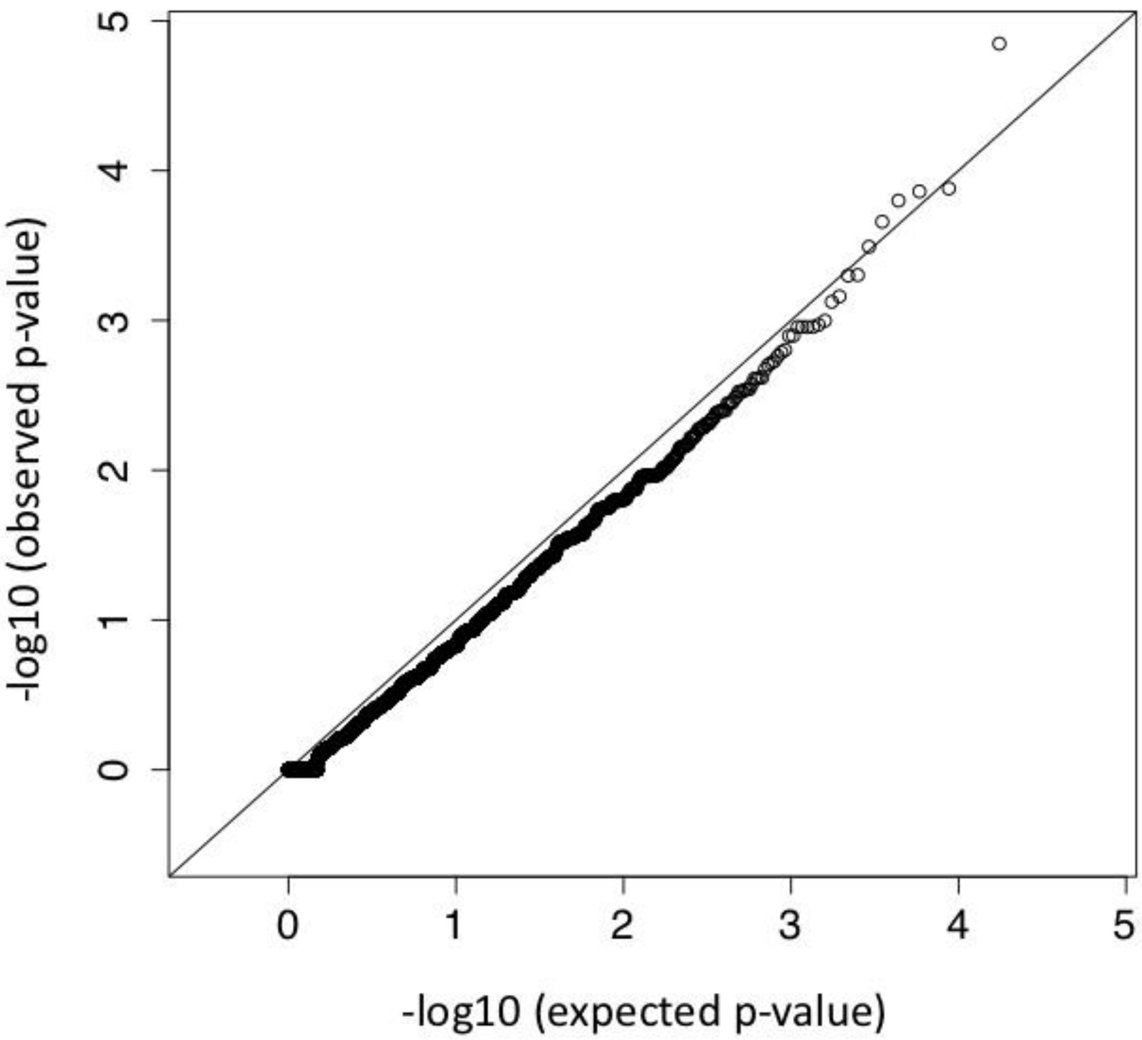

**Supplementary Figure 4**

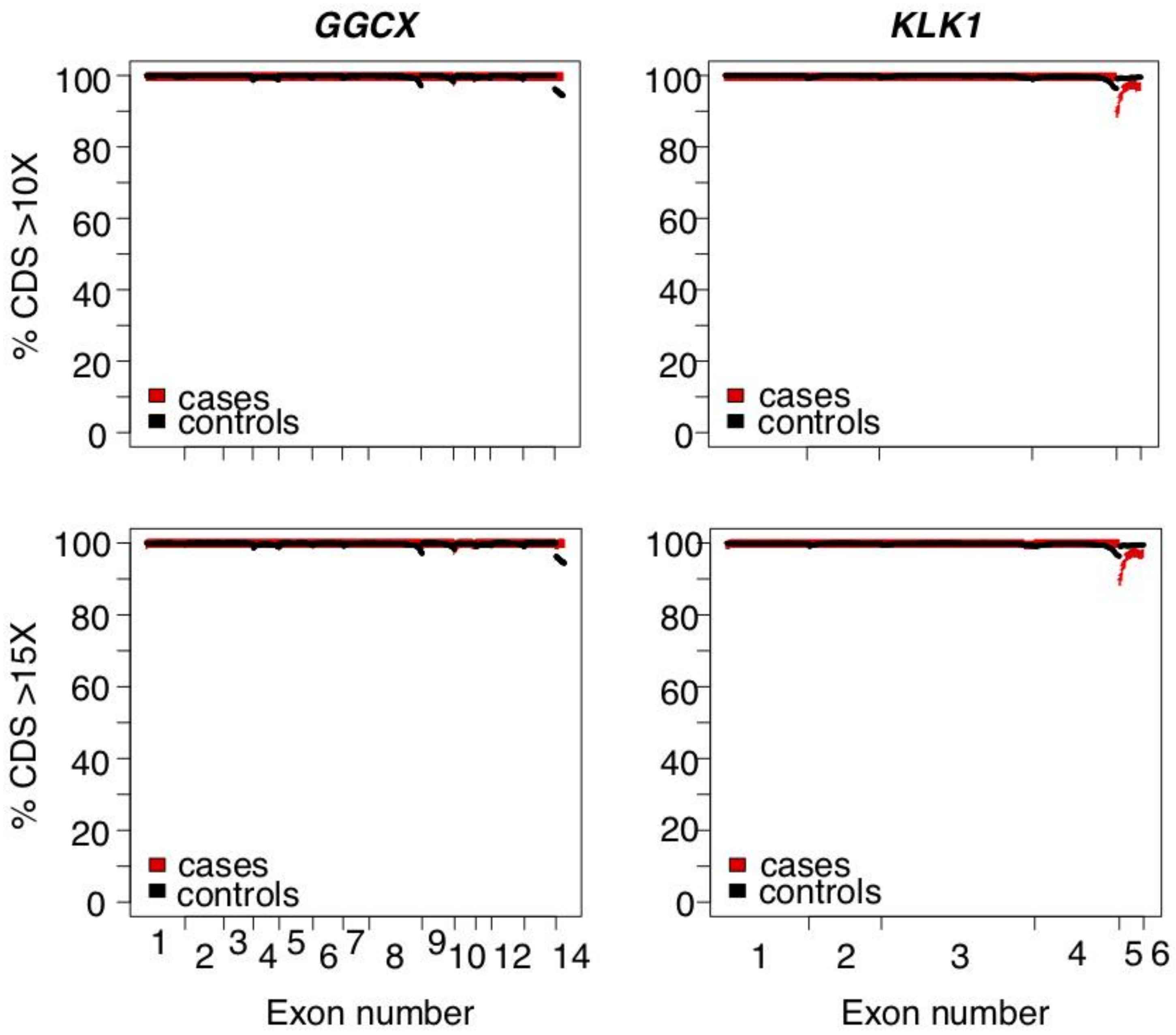
